# p66Shc is an apoptotic rheostat whose targeted ROS inhibition improves MI outcomes

**DOI:** 10.1101/2022.04.14.487897

**Authors:** Landon Haslem, Jennifer M. Hays, Hannah Schmitz, Satoshi Matsuzaki, Virginie Sjoelund, Stephanie D. Byrum, Kenneth M. Humphries, J. Kimble Frazer, Borries Demeler, Doris M. Benbrook, Ryan M. Tierney, Kelli D. Duggan, Franklin A. Hays

## Abstract

p66Shc is an oxidoreductase that responds to cell stress by translocating to mitochondria, where p66Shc produces pro-apoptotic reactive oxygen species (ROS). This study identifies ROS-active p66Shc as a monomer that produces superoxide anion independent of metal ions, inhibits cytochrome c peroxidase, and is regulated by environmental condition-induced structural changes. p66Shc anti-apoptotic functions, including: cytochrome c reduction, increased electron transport chain activity, and caspase cascade inhibition were also discovered. This study also demonstrates that p66Shc is a stress-dependent rheostat of apoptosis, regulated by p66Shc-mortalin complexes. These complexes decrease pro-apoptotic ROS production, without blocking p66Shc-mediated cytochrome c reduction. However, stress disrupts p66Shc-mortalin interactions, promoting apoptosis. Tipping p66Shc’s apoptotic balance toward anti-apoptotic functions by genetic knockdown or p66Shc-selective ROS inhibition decreased pro-apoptotic effects and improved outcomes in zebrafish myocardial infarction models, representing a potential new myocardial infarction treatment with promising results.

## INTRODUCTION

p66Shc is a ShcA (Src homology and collagen homology A) family member. The other members, p52Shc and p46Shc, share an amino acid sequence with p66Shc, but differ in length. All ShcA proteins contain identical PTB, CH1, and SH2 domains, but only p66Shc has a full N-terminal CH2 domain (CH2). p52Shc’s CH2 is truncated, and CH2 is absent in p46Shc (Pelicci et al., 1992). Since ShcA members only differ by CH2 length, functional differences are attributed to CH2 effects. ShcA proteins act as adaptor molecules that facilitate tyrosine kinase signaling complex formation (Ravichandran, 2001). This function is affected by CH2 length as p46Shc and p52Shc generally activate signaling pathways such as RAS/MAPK and mTOR pathways, while p66Shc’s effects are usually inhibitory (Gertz et al., 2008; Migliaccio et al., 1997; Soliman et al., 2014; Stevenson and Frackelton, 1998).

Another CH2-mediated difference is apoptosis regulation via oxidoreductase activity, which is only reported in the p66Shc isoform (Mir et al., 2020). Absent of stress signals, p66Shc localizes to mitochondria (mito, 44%), endoplasmic reticulum (24%), or cytoplasm as an inactive p66Shc-peroxiredoxin1 (Prx1) complex (32%) (Gertz et al., 2009; Orsini et al., 2004). Prevailing p66Shc ROS models suggest various cell stressors and signals increase cytosolic p66Shc phosphorylation at CH2 S36, primarily via PKCβ, JNK, or ERK and cause p66Shc mito intermembrane space (IMS) translocation (Hu et al., 2005; Le et al., 2001; Wills and Jones, 2012). Within the IMS, CH2-mediated p66Shc oligomerization can cause copper-dependent cytochrome c (cyt c) oxidation, producing H2O2 and triggering apoptosis (Gertz *et al*., 2008; Giorgio et al., 2005; Migliaccio et al., 1999). Yet, little is known about how p66Shc produces reactive oxygen species (ROS) or what factors influence ROS activity, and publications conflict about which form of ROS p66Shc produces (Mishra et al., 2019; Spescha et al., 2014; Vashistha et al., 2012). In reports using purified mouse CH2, Cys59-mediated tetramerization is required for ROS activity (Gertz *et al*., 2008) but other studies indicate p66Shc-cyt c interactions favor p66Shc monomers (Orsini *et al*., 2004; Orsini et al., 2006). Collectively, these studies indicate that Cys59-mediated p66Shc functions and Prx1 interactions inhibit ROS output in low stress, but high stress increases p66Shc activity and adverse pathological effects, which may require Cys59-mediated p66Shc tetramerization (Bugger and Pfeil, 2020; Carpi et al., 2009; Di Lisa et al., 2009; Kaludercic and Di Lisa, 2020; Wang et al., 2015; Zaccagnini et al., 2004).

Although necessary to maintain normal cell function, when [ROS] crosses a threshold, many pathologies are exacerbated. This includes ischemia/reperfusion injuries (IRI) such as myocardial infarction (MI) and stroke (Bugger and Pfeil, 2020; Incalza et al., 2018; Kaneto et al., 2010; Liguori et al., 2018; Mishra *et al*., 2019). Thus, ROS inhibition is a potential IRI therapeutic target (Cano Sanchez et al., 2018; Milkovic et al., 2019). Yet, global antioxidant administration is not effective in IRI clinical settings and can be harmful to recipients as it diminishes physiological ROS (Casas et al., 2020; Daiber et al., 2017; Kornfeld et al., 2015; Schmidt et al., 2015). These findings suggest targeting specific mito ROS-producers, contributing to excessive IRI [ROS], may improve MI outcomes by preventing pathological [ROS] thresholds (Andreadou et al., 2021; Casas et al., 2015; Di Lisa et al., 2017; Kornfeld *et al*., 2015). p66Shc fits these criteria as it is activated by IRI and increases mito [ROS], leading to apoptosis and poor outcomes (Baysa et al., 2015; Carpi *et al*., 2009; Di Lisa *et al*., 2017; Yang et al., 2014). In fact, p66Shc acts as a coronary artery disease biomarker and a stroke severity determinant (Franzeck et al., 2012; Noda et al., 2010; Spescha et al., 2015). Therefore, selectively inhibiting p66Shc-mediated ROS activity may provide substantial MI benefits.

Most primary MIs are not fatal in current hospital settings (Kontos et al., 2014). Yet, primary treatments (Tx) inadequately inhibit intrinsic cellular responses (*e*.*g*., ROS production). This forces non-infarcted cells to compensate for infarcted tissue, causing heart remodeling, heart failure (HF), and a 3-fold increase in post-MI (PMI) mortality over non-MI patients (Bugger and Pfeil, 2020; Gerber et al., 2016; Sutton and Sharpe, 2000). Since MI damage is proportionate to remodeling magnitude, decreasing initial MI damage by targeting excessive p66Shc ROS activity represents a potential remodeling prevention strategy that may mitigate MI damage, PMI remodeling, and HF (Baysa *et al*., 2015; Boengler et al., 2019; Di Lisa *et al*., 2017).

This study tests whether p66Shc’s apoptosis regulation can be manipulated to improve IRI outcomes by selectively targeting p66Shc ROS production. ROS production was inhibited using small molecule p66Shc-selective ROS inhibitors (identified in this study) in wildtype (WT) zebrafish and generating a p66Shc knockdown (KD) zebrafish line, then testing their effects in a cryoinjury MI model. A combination of *in vitro* and *in silico* approaches further characterized p66Shc mito functions, ROS mechanistic features, and structure-function relationships. This study unifies prior reports, using reported and novel functions, to demonstrate p66Shc is an apoptotic rheostat, which can be manipulated to improve MI outcomes.

## RESULTS

### p66Shc knockdown improves IRI outcomes in a ROS-dependent manner

Effects of targeted p66Shc inhibition on MI were explored with heterozygous p66Shc splice site zebrafish mutants expressing myocardial-specific eGFP (*myl7:eGFP:p66Shc*^*-/+*^,”KD” fish). KD and WT (*myl7:eGFP)* fish were cryoinjured to assess tissue injury and IRI outcomes (**Figure 1A**). For concision, all *in vivo* comparisons below are relative to corresponding positive controls and are significant (p-value < 0.05), unless otherwise noted. Resected hearts showed differences in wound responses via stereomicroscopy, tissue analysis, and untargeted proteomics (**Figure 1B-C**). KD fish reduced superficial injury size, trichrome fibrosis staining, and injury intensity by 61.4%, 65.2%, and 52.4%, respectively (**Figure 1B, S1A**). These data reflect attenuated injury expansion and are consistent with p66Shc KO mice having decreased PMI myocardial rupture (Baysa *et al*., 2015). We predicted differences were ROS-mediated as previous studies showed IRIs increase [ROS] and decreased [ROS] improves MI outcomes (Boengler *et al*., 2019; Di Lisa *et al*., 2017; Franzeck *et al*., 2012; Spescha *et al*., 2015). Indeed, WT injury site ROS fluorescence was 3-fold larger than KD (**Figure 1B**). Similar levels were observed in non-injured tissue and across entire heart sections (**Figure S1**). Scanning electron microscope (SEM) imaging showed regional cell recruitment and cell activation differences with high-activity regions proximal to cryoinjury sites and low-activity areas distal to active regions. In KD fish, active and total injury areas decreased by 47.6% and 29.7%, respectively (**Figures 1B, S2C**). Individually counting cells and comparing their morphology to active cells revealed decreases in KD fish total thrombocytes (66.2%), fibroblasts (72.2%), erythrocytes (53.9%) and leukocytes (45.8%) but showed non-significant increases in non-active fibroblast and thrombocyte counts. p66Shc KD also reduced scar tissue (58.9%), active injury (25.4%), and total injury (29.5%) (**Figures 1B, S2A-B**). In addition, KD fish showed organismal benefits, blocking cachexic effects and increasing PMI physical activity (**Figure S2D**).

**Figure 1.**
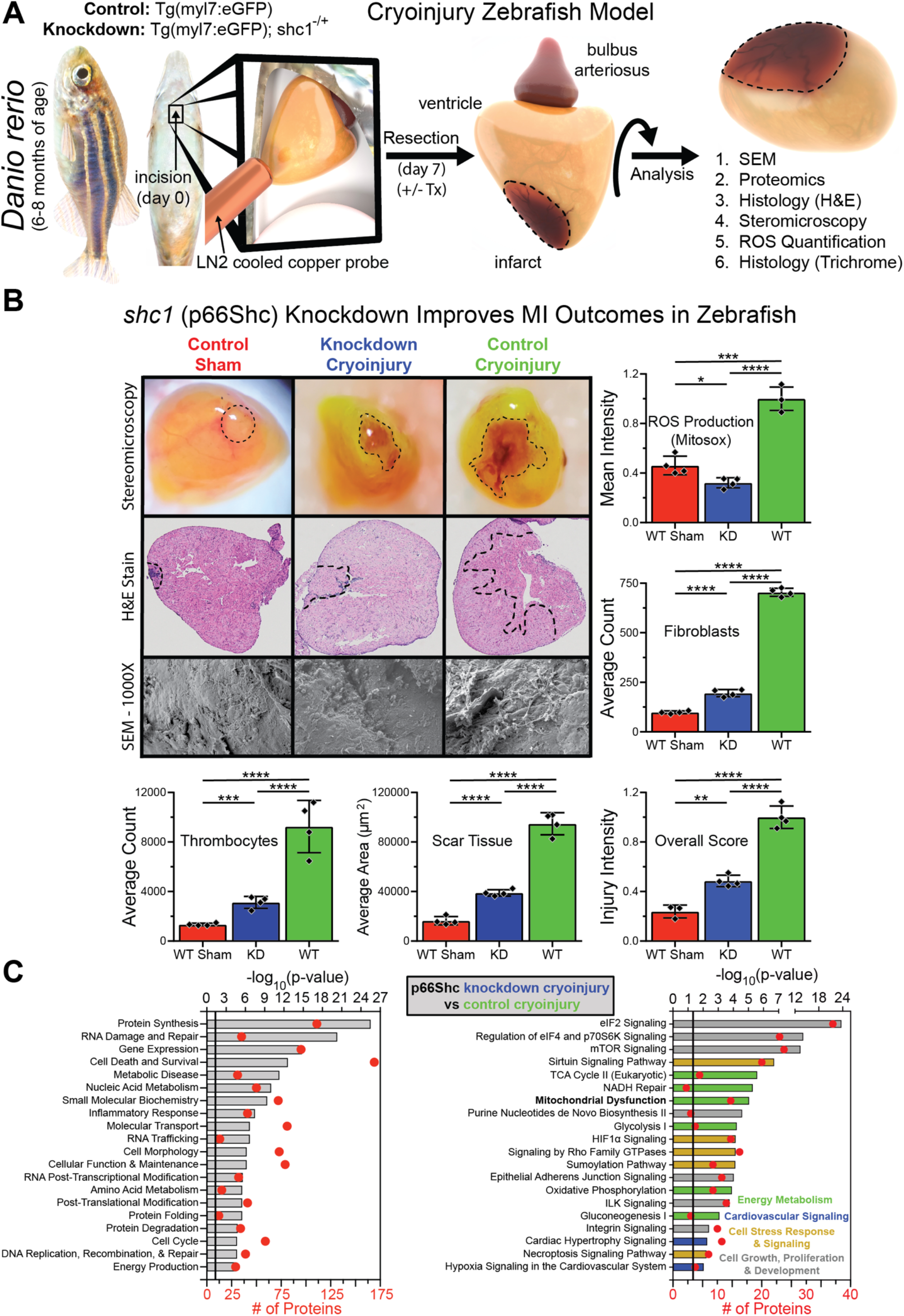
Shc1 (p66Shc) knockdown improves IRI outcomes in a ROS-dependent manner. (A) Cryoinjury zebrafish model. (B) Stereomicroscopy, H&E stains, and SEM representative MI images of in sham, KD, and positive control groups. Accompanying histograms illustrate image quantifications. Significance determined via t-tests. N=4. Data shown as mean ± SD. (C) PMI proteome analysis between KD and WT fish hearts. Groupings identified via Ingenuity Pathway Analysis (IPA, Qiagen) software. Shown as mean –log_10_ values and number of proteins in each group. N=3, significance at p-values :. 0.05.

PMI KD and WT fish ventricle proteome analysis provided protein-level insight into improved outcomes (**Figure 1C, Table S1**). Significant differences were observed in: protein synthesis, RNA damage and repair, cell death, inflammatory response, morphological alteration, protein degradation, and energy production. These processes showed altered signaling responses in four categories 1) energy metabolism, 2) cardiovascular signaling, 3) cell stress response signaling, and 4) cell growth, proliferation, and development. Other studies have identified similar metabolic pathway alterations in p66Shc KO mice (Hagopian et al., 2016; Hagopian et al., 2015). Expression profiles of differentially regulated proteins and pathways indicate that eIF2, mTOR, and redox (*e*.*g*., HIF1α) signaling are key mediators of p66Shc action in MI responses. In part, this response involves TCA cycle alterations, NADH repair, mito dysfunction, glycolysis, oxidative phosphorylation, and cardiac hypertrophy/hypoxia signaling. Relative to WT fish, PMI in KD fish have decreases in: [ROS], fibroblast and thrombocyte activity, infarct size, inflammatory cell recruitment, altered cardiac morphology, and decreases in: PMI body mass and physical activity (**Figure 1B-C, Figure S1-S2**).

### Small molecule p66Shc ROS inhibition improves IRI outcomes in a ROS-dependent manner

Four compounds that bind and inhibit human p66Shc *in vitro* ROS production, identified via functional screens, were selected for molecular-focused p66Shc cryoinjury functional studies: idebenone (Ide), carvedilol (Car), rapamycin (Rap), and N-acetyl cysteine (NAC) (**Figure 2A, Methods**). Ide is a coenzyme Q mimetic with antioxidant properties that improves insulin sensitivity in a ShcA-dependent manner (Jiang et al., 2021; Tomilov et al., 2018). Its mito mechanism is unknown (Buyse et al., 2003; Lyseng-Williamson, 2016). Car is a non-selective adrenergic blocker indicated in human heart failure and left ventricular dysfunction. Car alleviates diabetic cardiomyopathy in rats and improves heart remodeling, but its remodeling mechanism is unclear (Zheng et al., 2019). Rap is a macrolide used to treat human host vs. graft disease and vascular malformations, and experimentally treats mouse mito disorders (*e*.*g*., Leigh syndrome). Rap improves IRIs by inhibiting mTOR, increasing autophagy (Hadley et al., 2019). NAC is a non-specific antioxidant that increases glutathione synthesis, treats acetaminophen overdoses (associated with increased [ROS]), and benefits animals in muscular dystrophy or ischemic brain injury models (Cieslik et al., 2018). Based on binding affinity and ROS inhibition, Ide and Car Txs test selective p66Shc ROS inhibition, but Car also affects adrenergic signaling. Rap therapy tests mTOR inhibition, without affecting p66Shc ROS production, and NAC represents non-specific antioxidant Tx. Txs affected all measured parameters to varying extents (**Figures 2A, 2B, S1-3**), but consistent trends indicate selective p66Shc ROS inhibition is more beneficial than mTOR inhibition, and non-specific antioxidant Tx is harmful.

**Figure 2.**
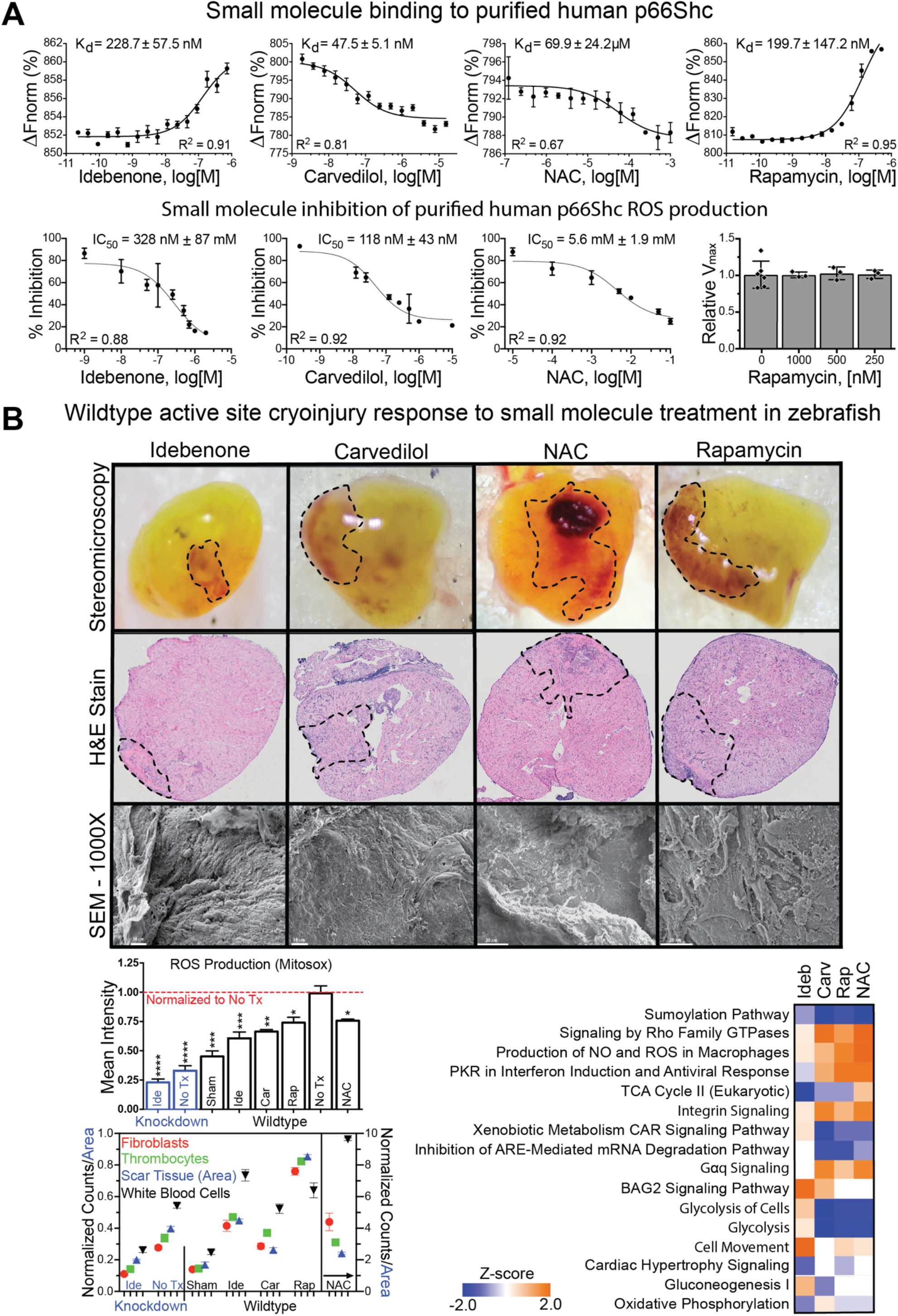
Small molecule p66Shc ROS inhibition improves IRI outcomes in a ROS-dependent manner. (A) MST binding and HE ROS experiments with the indicated compounds. K_d_ and IC_50_ values determined using non-linear regression. R^2^ values indicated. Shown as mean ± SD. N ≥ 3. (B) WT zebrafish PMI active site responses to small molecule Tx. Cell counts and intensity analysis performed as in **Figure 1B** with 3-4 hearts per group. N=3-4, and shown as mean ± SD, normalized to positive controls (WT No Tx). All cell counts were significant (P < 0.05), relative to controls. (C) Proteome changes illustrated as a Z-score heatmap. Blue indicates expression decreases, orange indicates increases. Shown as p-values vs # of identified proteins. N=3.

Ide, Car, and Rap Tx decreased the following parameters, respectively: injury intensity (52.4%, 33.5%, 17.0%), injury expansion/area (66.4%, 60.4%, 30.6%), fibrosis (67.0%, 55.1%, 25.7%), active injury area (26.8%, 63.5%, 24.9%), and MitoSox intensity (39.5%, 32.7%, 25.0%). Except for MitoSox intensity reduction (23.3%) NAC did not affect the above parameters and negatively affected many others (**Figure 2B, S1-2**). Non-active injury size differences between Tx groups were insignificant but treated hearts had reductions in all measured cell types except for NAC Tx, which increased cell counts (**Figure S2**). WT fish proteome analysis showed that Ide Tx caused decreases in TCA cycle, cardiac hypertrophy signaling, and oxidative phosphorylation (**Figure 2C, Table S1**). As an overall indicator of organismal PMI health, cachexic effects (body mass ratios) and physical activity (average distances) were measured. Ide increased PMI body mass by 4.8%, Car Tx had no effect, but Rap and NAC Tx decreased mass by 6.1 and 6.8%, respectively. Although, only Ide protected against cachexic effects, all Txs increased PMI physical activity (**Figure S2**). As a whole, results suggest that non-specific antioxidant administration worsens IRI prognoses, mTOR inhibition offers slight improvements, and selective p66Shc ROS inhibition provides substantial IRI protection.

### ROS-active p66Shc is a monomer that produces superoxide independent of metal ions

p66Shc’s prevailing ROS activity model suggests CH2 (**Figure 3A**) drives p66Shc oligomerization, causing copper-dependent H_2_O_2_ production and cyt c oxidation in the IMS (Giorgio *et al*., 2005; Mishra *et al*., 2019). Yet, precise mechanistic data is undetermined. We overexpressed, and purified, full-length human p66Shc and CH2 (**Figure S3E, S5A**) to characterize molecular function and mechanism. We demonstrate CH2 is intrinsically disordered with high conformational variability (**Figure 3A, 3D, S3A-B**). In addition, cyt c-free hydroethidine (HE) and electron paramagnetic resonance (EPR) spin-trapping assays demonstrate p66Shc, and CH2, produce superoxide anion (O_2_^-^) independent of cyt c electron (*e*^-^) transfer. However, O_2_^-^ activity was abrogated by superoxide dismutase (SOD) or hypoxia (**Figure 3B-C**).

**Figure 3.**
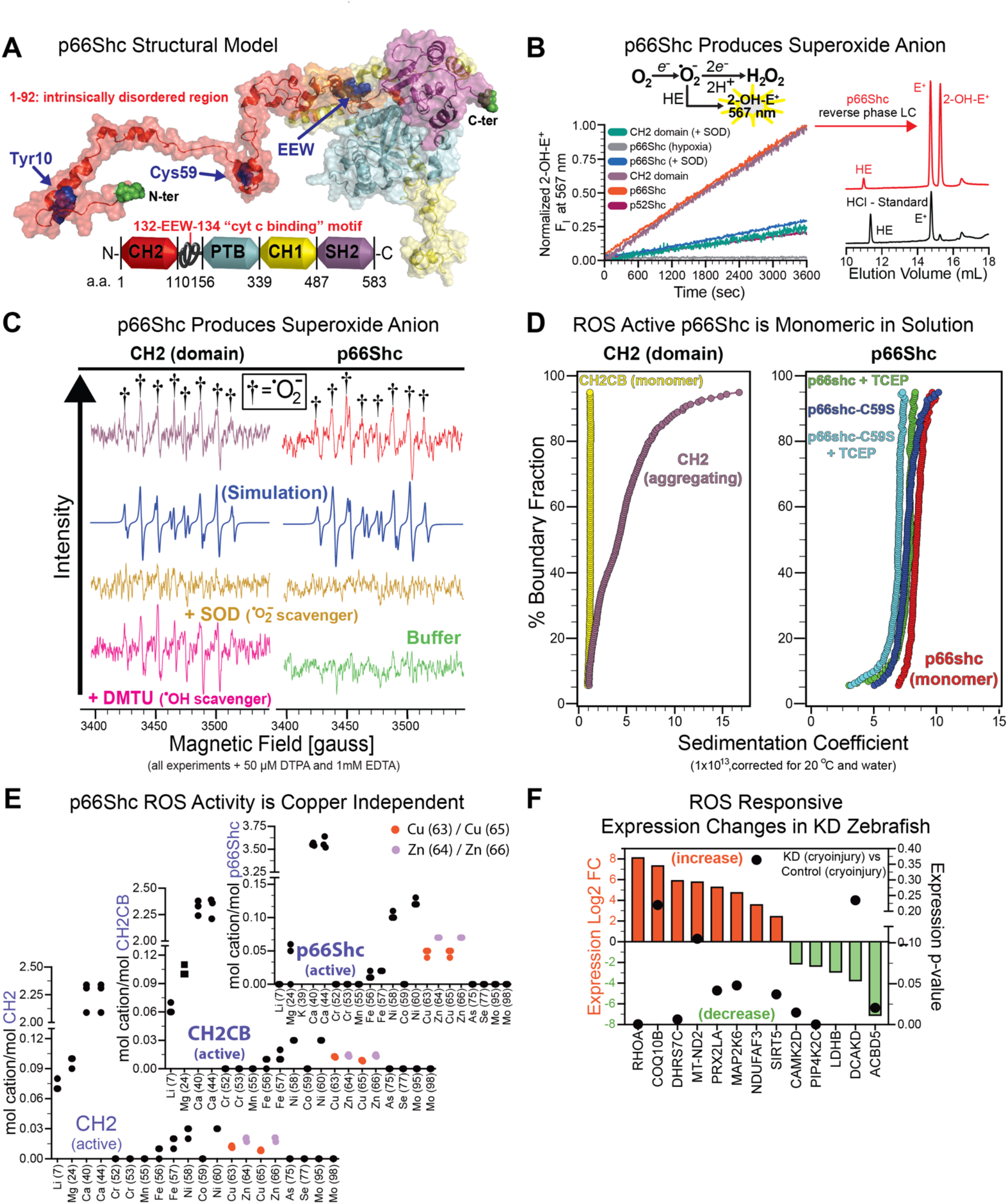
ROS-active p66Shc is a monomer that produces O_2_^-^independent of metal ions. (A) Human p66Shc structural model, colored by domain. Mutation sites for subsequent figures indicated in blue. (B-C) O_2_^-^ production assays via HE (B) and EPR (C). Shown as mean normalized fluorescence intensity (B) or signal intensity (C). N≥3 (unique p66Shc preparations). (D) AUC studies with CH2 and p66Shc. (E) ROS active ICP-MS analysis. Shown as means. N=3 (multiple unique preparations). Full statistics are available in **Table S2**. (F) PMI proteome analysis of increased (red) and decreased (green) ROS responsive protein expression between KD and WT fish. Data shown as expression log2(fold-change) and p-values (circles). N=3, significance at log values > 2 or p-values :. 0.05.

Furthermore, p66Shc wildtype is monomeric in solution with a molar mass in close agreement with mass spectrometry (MS) results (**Table S2**), and a frictional ratio of 1.8, which is indicative of an extended conformation (**Figure 3D**). Under reducing conditions, WT p66Shc’s sedimentation coefficient decreased, consistent with unfolding protein. The C59S mutation further enhanced this trend (**Figure 3D**). Likewise, the CH2CB (**Figure S3A**) molar mass was determined to be 14.7 kDa, consistent with a monomeric state. Purified CH2 was prone to aggregation (**Figure 3D**, left panel). Previous CH2 studies suggested C59 may drive higher order oligomerization (Galimov, 2010; Gertz *et al*., 2008). However, neither C59S mutation nor reductant disrupted O_2_^-^ activity or altered monomeric status (**Figure 3D, 5C**). Yet, reductant decreased p66Shc sedimentation coefficients by ∼0.3S (**Figure 3D**). This suggests that some of p66Shc’s 9 cysteine residues form intramolecular disulfide bonds. When reduced, p66Shc unfolds, increasing friction and solvent exposure. CH2 residues 1-110 are polydisperse in solution (**Figure 3D**), but extending constructs to include the putative “cyt c binding domain” (CBD) (Gertz *et al*., 2008) (**Figure 3A, S3A**) returns CH2 to a monomeric state (“CH2CB” in **Figure 3D**). These data suggest distal CH2 amino acids modulate conformational dynamics (supported by MD simulations in **Figure 6D**). Contrary to previous reports, we observed O_2_^-^ activity in copper-free assays with added metal chelators (**Figure 3B-C, S4A**). Also, inductively coupled plasma MS (ICP-MS) of active protein, purified without metal chelators, did not identify bound copper or zinc (**Figure 3E**).

Finally, proteomics data support KD fish had lower PMI oxidative and cellular stress, relative to WT fish (**Figure 3F, Table S1**). KD fish expression profiles were consistent with, but did not directly measure, decreases in: O_2_^-^ levels, oxidative stress, PIP5Kψ activity, peroxisome membrane contact sites and elongation, mito ROS, apoptosis, and increases in: GSH/GSSG ratio, oxidative stress capacity, reduced cellular environment, I/R cardioprotection, and PMI heart function (Aydin et al., 2019; Perez-Gomez et al., 2020; Ramadasan-Nair et al., 2014). These changes are further supported by observed improvements in PMI parameters noted in **Figures 1, S1-S2**.

### p66Shc regulates cyt c function and inhibits the caspase cascade

Since p66Shc produced O_2_^-^ independent of mitos or cyt c *e*^-^ transfer (**Figure 3B-C**), we investigated p66Shc-cyt c interactions and their effects. Microscale thermophoresis (MST) and surface plasmon resonance (SPR) measured human p66Shc-cyt c binding (**Figure 4A**). Results demonstrate p66Shc binds: ferrous cyt c (Fe^2+^, K_d_ = 15.8 μM), ferric cyt c (Fe^3+^, K_d_ = 38.5 μM), and K72A cyt c (peroxidase active, K_d_ = 29.7 μM) (Nold et al., 2017). Yet, cyt c is conformationally variable depending on lipid (*e*.*g*., cardiolipin (CL)) interactions and heme redox state (Patriarca et al., 2012). We assessed conformation effects with characterized cyt c mutants: H26Y (globular structure, high peroxidase activity) and Y67H (stabilized reduced state). H26Y mutants had markedly decreased p66Shc affinity (**Figure 4A**). p66Shc and cyt c reduction potentials are -35 mV and 17-250 mV, respectively (Battistuzzi et al., 2002; Giorgio *et al*., 2005). These potentials, and our observed binding affinities, thermodynamically support p66Shc-cyt c binding causes cyt c reduction, not oxidation. Indeed, co-incubating p66Shc or CH2 with cyt c (either Fe^3+^ or Fe^2+^) produces reduced cyt c and occurs in conditions limiting O_2_^-^ production (**Figure 4B, S5B**). For example, p46Shc lacks CH2, but reduces cyt c (**Figure 4B**), and Ide inhibits p66Shc O_2_^-^ activity while increasing p66Shc-mediated cyt c reduction 4-fold (**Figures 2A, S4B**). Results therefore support that cyt c reduction and O_2_^-^ production are independent.

**Figure 4.**
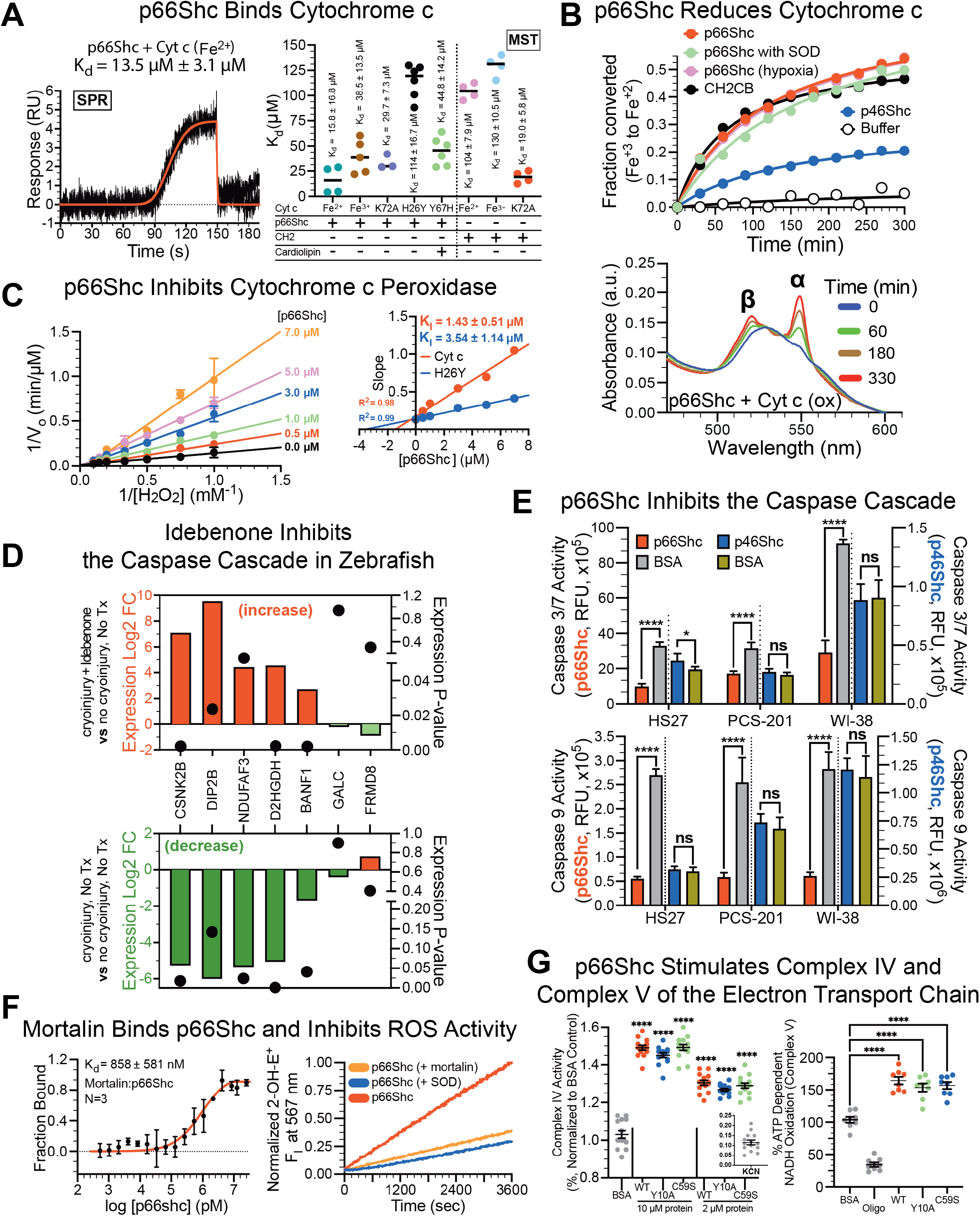
p66Shc regulates cyt c function and inhibits the caspase cascade. (A) SPR and MST p66Shc-cyt c binding results. SPR shown as means and trend line overlay. N=3 (2 protein preparations). MST shown as individual values. N ≥ 3 (multiple protein preparations). K_d_s shown as mean ± SD. (B) Oxidized (Fe^3+^) cyt c reduction by p66Shc, p66Shc + SOD, p66Shc + hypoxia (under p66Shc trace), and CH2CB. A550/521 absorbance ratios (bottom chromatogram) determined Fe^3+^-Fe2^+^conversion. Data shown as means. N = 3. (C) p66Shc-mediated cyt c peroxidase inhibition. Various [p66Shc] were tested against CL-activated cyt c or cyt c H26Y, using guaiacol and H_2_O_2_. Tetra-guaiacol absorption measured activity. Shown as mean ± SEM. N=3. (D and E) Proteome comparisons (D) of Ide-treated vs sham, and untreated vs sham fish. Data presentation and analysis as in **Figure 3F**. (E) Caspase 3/7 and 9 activity in human fibroblast cells incubated with p66Shc (orange), p46Shc (blue), or BSA (teal). Significance determined via t-tests. Shown as mean ± SD. N=5. (F) p66Shc-mortalin MST binding results and effects on ROS activity. K_d_ determined by non-linear regression. Shown as mean ± SD. N=4. (G) Complex IV and V, activity assays. WT, Y10A, or C59S p66Shc was incubated with fresh mitoplasts followed by Complex IV or V activity assays. Data shown as normalized mean ± SEM and individual points. Complex IV N=14, Complex V N=8. Significance determined via t-tests.

**Figure 5.**
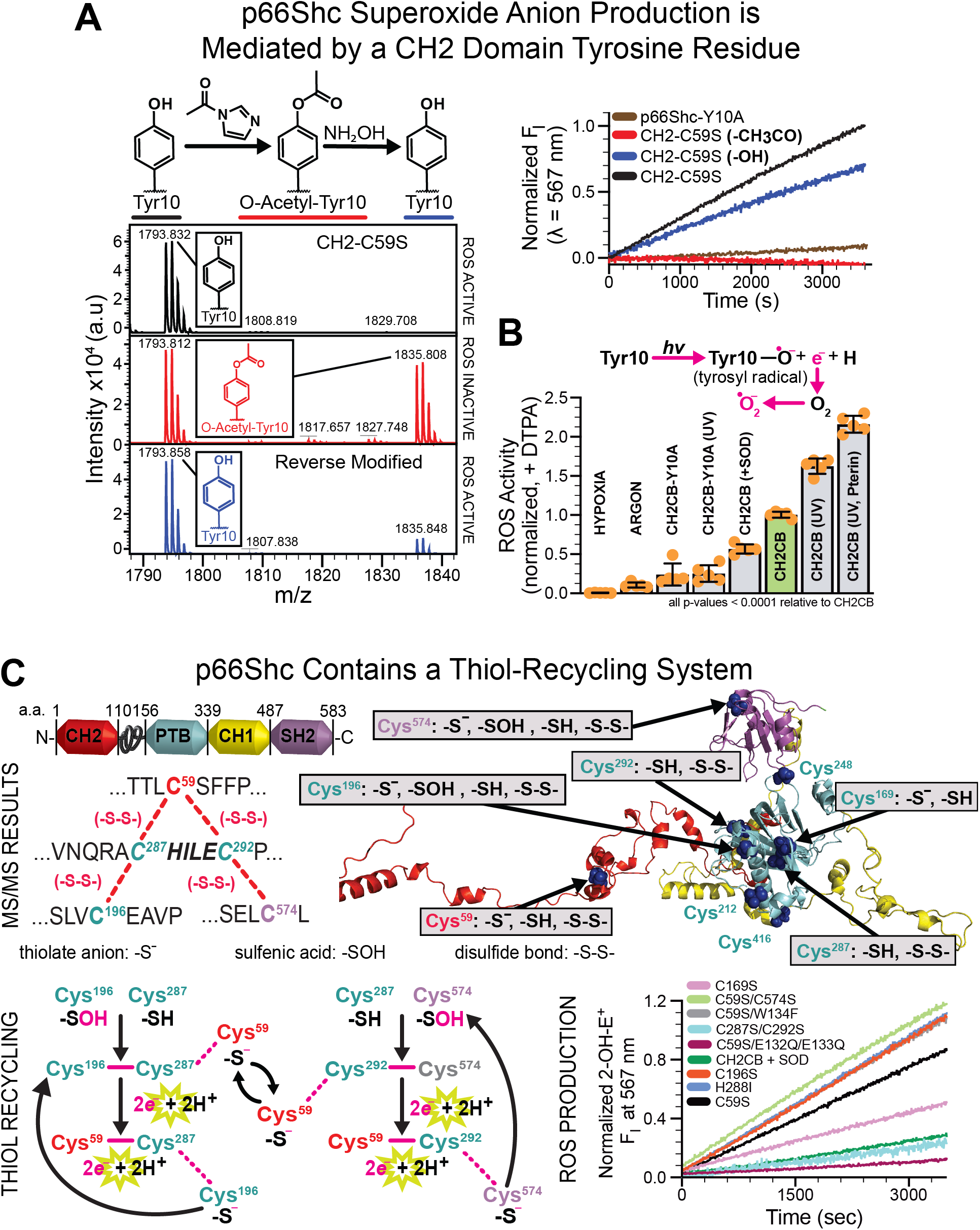
p66Shc utilizes a thiol-recycling system that facilitates O_2_^-^ production in a Y10- and O_2_-mediated reaction. (A) MS and HE analysis of modifications that modify ROS activity. (A) Unmodified CH2 (black) loses ROS activity upon acetylation (red) which is regained after deacetylation (blue). Y10A mutation abolishes ROS activity (brown trace). F_I_ traces shown as means. N=3 (unique protein preparations). HE studies (histogram) testing inhibitory and enhancing ROS conditions. Shown as mean ± SEM normalized to CH2CB (green bar) and individual points. All comparisons to CH2CB, using t-tests are significant (p-values <0.0001). N=5. (B) MS Cys interaction mapping in human p66Shc. Interactions indicated in structural and conceptual p66Shc models, colored by domain. Mutagenesis studies show inhibited ROS activity with a C287/C292 double mutant (light blue trace). HE assays presented as normalized means. N=3.

**Figure 6.**
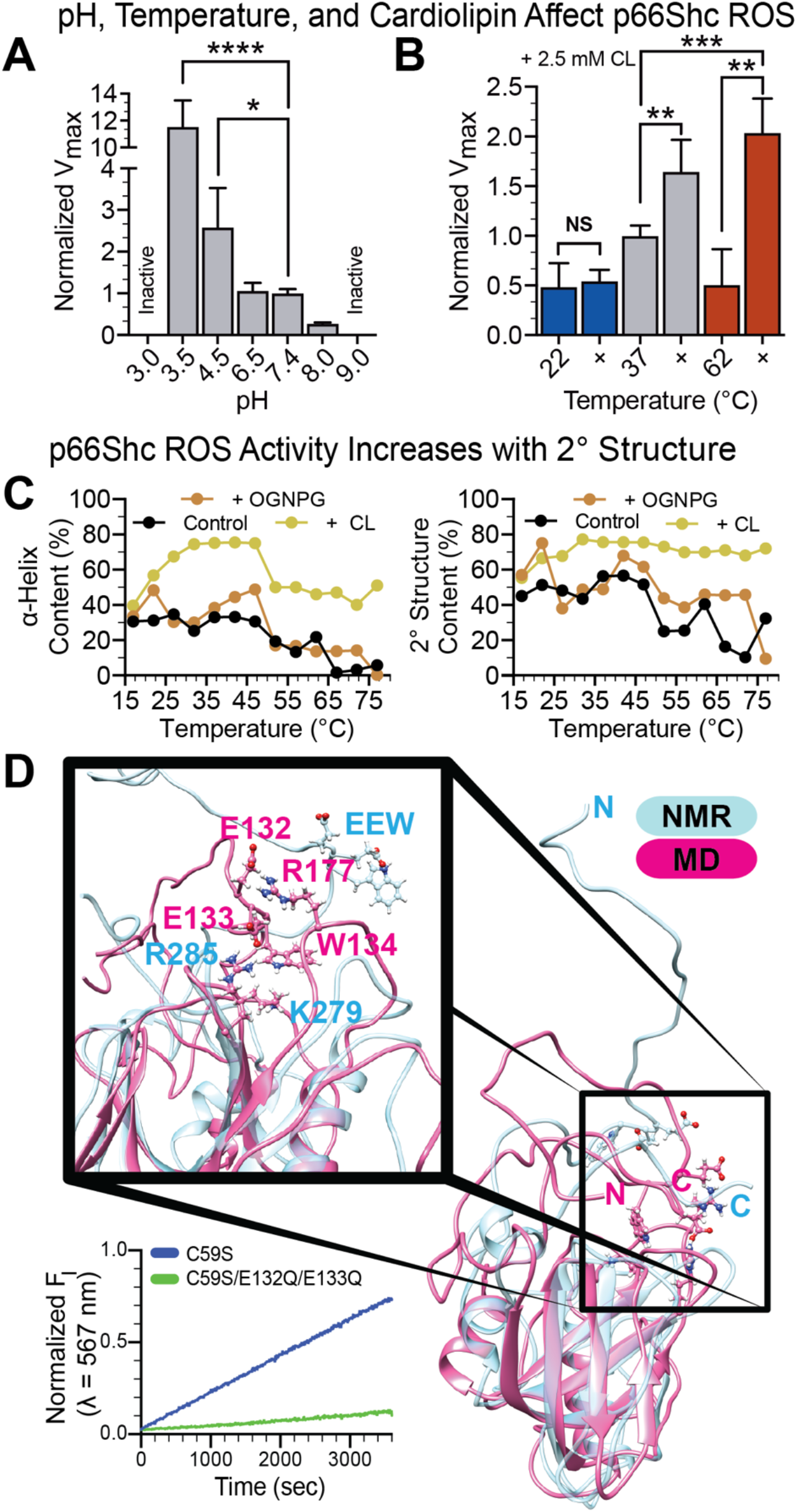
IRI stress responses promote p66Shc ROS-active conformational changes and increase O_2_^-^ production. (A-B) p66Shc HE activity assays with increasing pH (A) or temperature (B) ± CL. V_max_ values determined using Michaelis-Menten non-linear regression analysis of 2-OH-E+ corrected HE velocity data. Shown as mean ± SEM. Significance determined via t-tests. N=3 independent measurements. (B-C) Human p66Shc activity increases in warm, lipid-rich, environments. (C) CH2 CD analysis. Data shown as predicted α-helix content or predicted total secondary structure vs temperature. Predictions were made using raw data means in DichroWeb’s fourth reference data set (CONTIN). N=3. (D) MD analysis (magenta) relative to a previous PTB domain NMR structure (cyan, 1N3H). Analysis identified putative cation-π and salt bridge interactions that stabilize an active p66Shc conformation. Removing these interactions via C59S/E132Q/E133Q mutagenesis abrogates activity. Shown as normalized means. N=3.

Since p66Shc binds peroxidase-active cyt c, a key lipid peroxidation and apoptotic response modulator (Kagan et al., 2009), we tested p66Shc’s effects on peroxidase activity. Results demonstrated non-competitive inhibition with a K_i_ of 1.43 μM (**Figure 4C**). H26Y cyt c (peroxidase-active) assays had a similar K_i_ of 3.54 μM, closely agreeing with H26Y-cyt c minocycline inhibition (K_i_ = 1.0 μM) (Patriarca *et al*., 2012). Furthermore, CH2 peroxidase assays decreased H_2_O_2_ breakdown by 48.9% relative to CH2-free assays (**Figure S4C**). Thus, observed [H_2_O_2_] increases in previous studies may be caused by p66Shc-mediated cyt c peroxidase inhibition and O_2_^-^ spontaneous dismutation. Ide Tx, which notably decreased myocardial [ROS], caused proteome changes consistent with caspase inhibition (Adlat et al., 2020; Cox et al., 2011; Li et al., 2002; Perez-Gomez *et al*., 2020; Sugiana et al., 2008) (**Figure 4D, Table S1**), indicating either [ROS] or p66Shc directs caspase activity in stressed environments.

p66Shc effects on caspase inhibition were therefore tested in human fibroblast lines (dermal: HS27 and PCS-201, lung: WI-38). p66Shc decreased caspase 9 activity ∼5-fold and caspase 3/7 activity ∼3-fold, but p46Shc (no CH2) did not affect caspase activity (**Figure 4E, S5C**). These data support that caspase cascade inhibition is CH2-dependent. Yet, it is unclear if inhibition is direct or indirect (*e*.*g*., O_2_^-^ production, cyt c sequestration). ShcA proteins bind a diverse array of cellular proteins to modulate signaling pathways (Bhat et al., 2015; Zheng et al., 2013). Thus, p66Shc-dependent caspase cascade regulation may involve binding interactions. Prior work suggests regulation occurs, in part, via *HSPA9* (mortalin) association (Kang et al., 2018; Liu et al., 2005; Londono et al., 2012; Orsini *et al*., 2004). In response to moderate oxidative stress, mortalin is upregulated and protects cells by preventing pro-apoptotic p53 transcription and forming a stress-dependent complex with p66Shc (Gestl and Anne Bottger, 2012; Lu et al., 2011; Orsini *et al*., 2004; Sadekova et al., 1997). MST validated mortalin-Bcl2 (K_d_ = 171 nM) and mortalin-p66Shc (K_d_ = 858 nM) binding interactions (**Figure S5E**). Mortalin inhibited p66Shc O_2_^-^ activity (**Figure 4F**) without impacting p66Shc-mediated cyt c reduction (not shown), suggesting that cell stress frees p66Shc from an anti-apoptotic complex and induces a switch to pro-apoptotic activity (Orsini *et al*., 2004). This is consistent with a previous observation that mortalin depletion induces cell death via oxidative stress and Bcl-2 regulation (Starenki et al., 2015).

p66Shc-mediated cyt c reduction has electron transport chain (ETC) implications. To test ETC effects, we assayed Complex IV and V activity with p66Shc and respiring C57B/6J mouse heart mitoplasts. Purified p66Shc (ROS active), p66Shc-Y10A (ROS inactive), and p66Shc-C59S each increased complex IV activity in a concentration-dependent manner (2 μM:∼30%; 10 μM:∼60%) (**Figure 4G**). Likewise, Complex V activity increases ∼50% for each protein construct. Assays were not affected by SOD (not shown).

### p66Shc utilizes a thiol-recycling system that facilitates O_2_^-^ production in a Y10- and O_2_-mediated reaction

Understanding p66Shc molecular function requires mechanistic data. Thus, we investigated how mutations affect ROS activity. p66Shc C59S did not affect oligomeric state, ROS production, or cyt c activities (**Figures 5A, S3D**). However, mutating CH2’s strictly conserved Y10 residue to Y10A prevented O_2_^-^ production (**Figure 5B, S3F**). Acetylating Y10’s – OH group also blocks O_2_^-^ activity but the block is reversed when deacetylated (**Figure 5A**), suggesting Y10 is critical in O_2_^-^ production. Tyr radical intermediates are important in many ROS-active enzymes and often have a 410 nm absorption peak, which was indicative of ROS activity in p66Shc and CH2 preparations (**Figure S5A**) (Blomberg, 2016; Lehrer and Fasman, 1967). UV exposure can form Tyr radicals (Leo et al., 2013; Stubbe and van Der Donk, 1998) and UV-exposed CH2 increased O_2_^-^ production by 57.3%. Assays including UV and pterin, a UV sensitizer, increased ROS activity > 2-fold. This suggests that O_2_^-^ generation may involve a Tyr radical (**Figure 5B**). Radical formation increases with O_2_ availability (Stubbe and van Der Donk, 1998). Limiting O_2_ availability via argon purge lowered O_2_^-^ production by 88.9% and hypoxia abrogated activity, further supporting radical involvement (**Figure 5B**).

We hypothesized that intramolecular disulfide bonds (**Figure 3D**) contributed to ROS activity and comprehensively mapped p66Shc cysteine interactions. ROS-active p66Shc MS mapping revealed an intramolecular thiol-recycling system with observed exchanges between disulfide (-S-S-), thiolate anion (-S^-^), and sulfenic acid (-SOH) intermediates (**Figure 5C, Table S2**). C287 and C292 are located in a conserved PTB domain motif (C287-HILE-C292-P) and each interacted with 2 cysteines (**Figure 5C**). C287 and C292 both disulfide bonded C59 but disulfide bonds at C196 and C574 were exclusive to C287 and C292, respectively. Dimedone, N-ethylmaleimide (NEM), and iodoacetamide (IAM) MS studies identified -SOH intermediates at C196 and C574 while C59, C169, C196, and C574 also existed as -S^-^ (**Figures 5C, Table S2)**. Mutation effects illustrated thiol recycling promotes O_2_^-^ generation via the C287/C292 dyad (**Figure 5C**). No individual mutation blocked ROS activity to a similar degree (*e*.*g*., C169S, C59S, C196S, C574) as the C287S/C292S double mutant, which had similar O_2_^-^ activity as Y10A mutants or WT p66Shc in SOD’s presence. Mutating the CBD (EEW, **Figure 3A**) either attenuates or fully disrupts ROS activity when combined with specific mutations (*e*.*g*., C59 or C287) (**Figure 5C**). A more detailed -EEW-motif analysis is provided below. Thus, MS data demonstrate complex interdomain chemistry and conformational changes that modulate O_2_^-^ production.

### IRI stress responses promote p66Shc ROS-active conformational changes and increase O_2_^-^ production

In addition to increased p66Shc-mediated [ROS] (**Figure 1**), IRIs also cause wound-site acidification, shear stress, and increased tissue temperature (Lagadic-Gossmann et al., 2004; Liu et al., 1998; Silvia, 2006; Yan and Kleber, 1992). Thus, a key question in IRI pathology is: do IRI effects alter p66Shc ROS activity? We answered this question by simulating IRI changes in pH, shear stress, temperature, and lipid environment. p66Shc reached maximal O_2_^-^ activity between a pH of 3.5 and 4.5 (**Figure 6A**). Activity also increases with shear, temperature, or CL (an IMM component) exposure (**Figure 6B, S5F) Table S1**). Thus, environment modulates p66Shc ROS production and IMS conditions significantly increase ROS activity. These results suggest the identified Cys network (**Figure 5C**) is rate-limited, and potentially regulated, by Cys leaving groups’ electrophile strength during interchanges (Nagy, 2013). Circular dichroism (CD) investigated p66Shc secondary structure content and found that conditions significantly increasing V_max_ increased ShcA α-helical content (**Figure 6C**). In addition, CL presence was key in post-thermal melt α-helix preservation, refolding, and ROS activity (**Figures 6B, S5D**). CL-induced α-helix increases were greater for CH2 than p66Shc, suggesting CL mediates O_2_^-^ production by stabilizing a CH2 ROS-active conformation.

The CBD may regulate environmental effects and O_2_^-^ activity as MD simulations suggest CBD coordinates CH2-PTB orientation (**Figure 6D**). In simulations, the CBD forms salt bridges and cation-π interactions with opposing PTB R177, R285, and K279 residues, respectively (**Figure 6D**). These interactions facilitate partial folding that orients the PTB N-terminus toward its core. With the previous NMR study (Farooq et al., 2003), these results indicate significant CBD structural fluctuations on a ns to ms timescale that can orient CH2 relative to PTB. E132Q/E133Q double mutant simulations lost important folding interactions (**Figure 6D**), causing the PTB N-terminus to orient away from its core. Simulation results were tested by disrupting simulated CBD salt bridges via p66Shc E132Q and E133Q mutations, causing a significant decrease in O_2_^-^ production (**Figure S6**). This supports a previous study where this mutant did not induce pro-apoptotic mito swelling (Giorgio *et al*., 2005). Thus, the CBD modulates ROS activity via salt bridge interactions that prime p66Shc’s ROS-active conformation, not via direct cyt c binding.

## DISCUSSION

ROS is a paradoxical cell function modulator. High [ROS] is harmful in diabetes, heart disease, and some cancers while moderate [ROS] is protective (*e*.*g*., hormesis, energy metabolism and cell signaling) (Antonucci et al., 2021; Calabrese and Mattson, 2017) but very low [ROS] is detrimental (Milkovic *et al*., 2019). p66Shc function is also paradoxical. For example, decreased p66Shc-mediated ROS production or expression in IRIs is beneficial (Baysa *et al*., 2015; Boengler *et al*., 2019; Mishra *et al*., 2019; Spescha *et al*., 2015). Yet, p66Shc KO increases short-term myocardial injury, negatively impacts ischemic preconditioning, and decreases lifespan in natural environments (Akhmedov et al., 2015; Giorgio et al., 2012; Heusch, 2015). Here, we define p66Shc mito function as a tunable rheostat that governs survival and pro-apoptotic responses in a stress-dependent manner, uniting and resolving conflicting findings where p66Shc is either beneficial or detrimental to human health and disease.

This study demonstrates monomeric p66Shc responds to environmental conditions to produce O_2_^-^, independent of cyt c or copper, with activity mediated by: cys-interactions, Y10 radical formation, and O_2_ availability (**Figures 3, 5, S3A)**. Identifying p66Shc’s ROS product as O_2_^-^ is consistent with reports where MI was associated with increased post-ischemic [O_2_^-^], ischemia severity, and O_2_^-^-dependent negative PMI remodeling (Kramer et al., 1987; Siwik et al., 1999; van Deel et al., 2008). These results may also explain, in part, why hypoxia is beneficial in diseases where decreased p66Shc-mediated ROS improves outcomes (*e*.*g*., Leigh syndrome, MI) (Jain et al., 2019; Pang et al., 2018; Wojtala et al., 2017).

Although CH2 is mostly disordered (**Figure 3A, S3A-C**), conditions increasing ROS activity also increase α-helix content, with optimal p66Shc O_2_^-^ production occurring in conditions that mimic MI: low pH, high temperature, high shear stress, and increased CL exposure (Boengler *et al*., 2019; Di Lisa *et al*., 2017; Gao et al., 2020; Kumar et al., 2018; Liu *et al*., 1998; Merrill et al., 2020; Spescha *et al*., 2014) (**Figure 6A-C, S5F**). Thus, our data suggest p66Shc senses environmental changes (*e*.*g*., pH) caused by IMS translocation, or cell stress, driving ROS active conformational switching via α-helix stabilization (**Figure 6**). p66Shc also increases [ROS] by inhibiting cyt c peroxidase activity (**Figures 4C, S4C**). Thus, observed mito H_2_O_2_ “accumulation”, identified in previous reports, occurs via O_2_^-^ spontaneous dismutation and cyt c peroxidase inhibition.

Our studies also provide ROS generation mechanistic insights. Sequence analysis identified a single conserved CH2 Tyr residue (Y10) and ROS experiments demonstrate Y10 is required for ROS formation (**Figure 5, S3F**). Furthermore, O_2_^-^ production requires a single *e*^*-*^ transfer and conditions that stimulate radical formation increased CH2 O_2_^-^ generation, suggesting potential Y10 radical formation. Tyrosine radicals are involved in many important biological reactions, including those catalyzed by cyt c oxidase, and as intermediates in *e*^-^ relays (Blomberg, 2016; Giese et al., 2009; Stubbe and van Der Donk, 1998). Y10 appears to be part of an *e*^-^ relay, as preventing *e*^-^ transfer via Y10A mutation, or reversible acetylation, disrupted O_2_^-^ production with minimal impact on cyt c interactions (**Figure 5A**). Such *e*^-^ relays are dependent on *e*^-^ sinks, a role that could be filled by the oxyanion hole identified through *ab initio* modeling (**Figure 6D**). Comprehensive p66Shc MS analysis (**Figure 5C, Table S2**) identified an interdomain thiol-recycling system capable of providing conformational variation required for Y10-oxyanion hole interactions. This system is also required for ROS activity (**Figure 5B**). MD simulations further support conformation-dependent ROS activity as losing key interactions, which maintain a simulated CH2-PTB orientation favoring *e*^-^ flow, prevented experimental ROS production (**Figure 6D**). We also hypothesize that conformational variability, and the thiol-recycling system, modulates protein-protein interactions within the IMS (Habich et al., 2019).

ROS assays were conducted without cyt c, demonstrating O_2_^-^ production is independent of p66Shc-cyt c-interactions. This raised important questions, including – does p66Shc interact directly with cyt c and, if so, what are the functional consequences? Indeed, p66Shc and cyt c bind (**Figure 4A**), but its binding affinity is lowest with the constitutively peroxidase active H26Y cyt c mutant (Patriarca *et al*., 2012). Thus, p66Shc may prevent cyt c peroxidase formation and pro-apoptotic signaling by binding, and stabilizing, non-peroxidase active cyt c conformations (Kagan *et al*., 2009). We also observed p66Shc-mediated cyt c reduction, with or without SOD (**Figure 4B**). In addition, Ide increases p66Shc-mediated cyt c reduction while inhibiting O_2_^-^ production – suggesting ROS activity and cyt c reduction are uncoupled. However, cyt c reduction may play a stress-dependent role in regulating mito protein composition since protein import is governed by cyt c redox status (Habich *et al*., 2019; Riemer et al., 2009; Stojanovski et al., 2008). Our *in* vivo results suggest protein composition changes, from Tx’s that enhance p66Shc-mediated cyt c reduction, are beneficial in IRI and improve adverse cardiac morphology changes proportionate to p66Shc-mediated cyt c reduction (**Figures 2, S1-2, S4B**). In addition, reduced cyt c is linked to protective IRI effects and increased ETC activity (Di Lisa *et al*., 2017; Kalpage et al., 2019) and we found that p66Shc increased ETC activity independent of ROS production. ETC enhancement may occur via cyt c recycling or direct *e*^-^ transfer to Complex IV but requires further studies to confirm. This is consistent with prior studies where p66Shc activation increased ETC activity and [ROS] while p66Shc KO decreased ETC efficiency (Berniakovich et al., 2008; Lone et al., 2018).

Further studies revealed that p66Shc can influence apoptosis via caspase cascade inhibition (**Figure 4D-E**). These findings are consistent with an earlier report, where p66Shc KO mice in IRI models had increased caspase 3-mediated apoptosis during short-term ischemic insults (Akhmedov *et al*., 2015). Yet, p66Shc did not inhibit purified caspase 3 activity (**Figure S5C**), suggesting inhibition occurs after IMM pore formation (D’Arcy, 2019). Mortalin is a stress sensor residing throughout the cell that can interact with p53, Bcl-2, p66Shc, and CL (Dores-Silva et al., 2020; Orsini *et al*., 2004; Ran et al., 2000; Saxena et al., 2013). Moderate oxidative stress upregulates mortalin, which protects cells by inhibiting pro-apoptotic p53 transcription and p66Shc ROS activity via cytoplasmic sequestration and formation of a stress-sensitive anti-apoptotic complex, respectively (Orsini *et al*., 2004) (**Figure 4F)**. Since high oxidative stress or UV stimulation frees p66Shc and causes mito trans-membrane potential collapse, cell stress may mediate a transition to pro-apoptotic p66Shc activity (Orsini *et al*., 2004).

KD fish had a ROS-dependent improvement in all measured parameters (**Figures 1, S1-S2**) and selectively inhibiting p66Shc ROS in WT fish via Ide or Car Tx showed improvements similar to KD fish in: injury intensity, PMI physical activity, fibrosis, histological injury expansion, SEM active injury area, and injury cell counts (**Figures 2, S1-S2**). Also, Ide treatment may inhibit zebrafish caspase cascades during MI (**Figure 4D**). These results suggest an adjusted p66Shc rheostat set-point improved outcomes. The importance of targeting p66Shc ROS production over general ROS inhibition is underscored by NAC results. Despite lowering [ROS], NAC was detrimental in nearly all parameters (**Figures 2, S1-S2**). Thus, our data support previous conclusions that selective ROS inhibition of mito proteins, activated by IRI, would provide substantial IRI improvements (Andreadou *et al*., 2021; Casas *et al*., 2015; Di Lisa *et al*., 2017; Kornfeld *et al*., 2015). Indeed, MI therapies lowering mito [ROS], such as coenzyme Q (CoQ), mitoquinone (mitoQ), mito-TEMPO and Bendavia, show pre-clinical or clinical PMI benefits. MitoQ and mito-TEMPO improve PMI cardiac contractility, heart dilation, and cardiac graft results by scavenging mito ROS and lowering pro-apoptotic cyt c release (Dey et al., 2018; Ni et al., 2016). Of note, Ide shares a pharmacophore with mitoQ and CoQ, suggesting their beneficial effects may be p66Shc-mediated. Bendavia decreases oxidative stress, infarct size, apoptosis, and detrimental heart remodeling while increasing ETC activity in animal models by binding CL, which prevents ROS-induced CL oxidation and cyt c peroxidase formation (Birk et al., 2013; Kloner et al., 2012). Since p66Shc also appears to prevent cyt c peroxidase formation, results similar to Bendavia’s may be possible via p66Shc modulation.

Our results combined with prior studies, support the following unified p66Shc model in low-, and high-, stress conditions (**Figure 7**). In low-stress (**Figure 7B**), cytoplasmic p66Shc is monomeric and primarily inactivated by environment, Prx1, or mortalin. In addition to p66Shc inhibition, mortalin inhibits p53-mediated apoptosis, and may stabilize Bcl-2 proteins in low stress (Gestl and Anne Bottger, 2012). A small portion of p66Shc is localized to mitos to produce basal O_2_^-^, stimulate ETC activity, and prevent cyt c lipid peroxidation. In contrast, high-stress (*e*.*g*., IRI; **Figure 7C**) disrupts mortalin complexes and increases p66Shc mito localization (Galimov, 2010; Orsini *et al*., 2004). Increased free mito [p66Shc] increases [ROS] and CL oxidation, leading to cyt c dissociation from the IMM that causes IMM CL depletion and subsequent ETC dysfunction (Covian and Trumpower, 2006; Paradies et al., 2004; Paradies et al., 1999; Petrosillo et al., 2003). Continued stress promotes p53-mediated p66Shc upregulation, which increases p66Shc-mediated Foxo3A sequestration and [ROS], preventing antioxidant transcription (Chahdi and Sorokin, 2008; Trinei et al., 2002). Unable to further pass *e*^-^s to a dysfunctional ETC, inhibit caspase activation, or prevent cyt c peroxidase formation, p66Shc activity becomes strictly pro-apoptotic. In addition, p66Shc becomes increasingly ROS active with IRI environmental changes (*e*.*g*., pH). These effects synergize to ultimately stimulate IMM pore formation and apoptosis. However, targeted p66Shc ROS inhibition can mitigate IRI outcomes by promoting p66Shc’s pro-survival functions. Therefore, tipping p66Shc’s apoptotic balance toward anti-apoptotic functions by p66Shc-selective ROS inhibition represents a potential new MI treatment with promising results. In contrast, disrupting p66shc-mortalin interactions with drugs, such as SHetA2, tip the balance toward pro-apoptotic functions; a concept under clinical investigation (NCT04928508, (Benbrook et al., 2013; Chandra et al., 2021; Ramraj et al., 2019).

**Figure 7.**
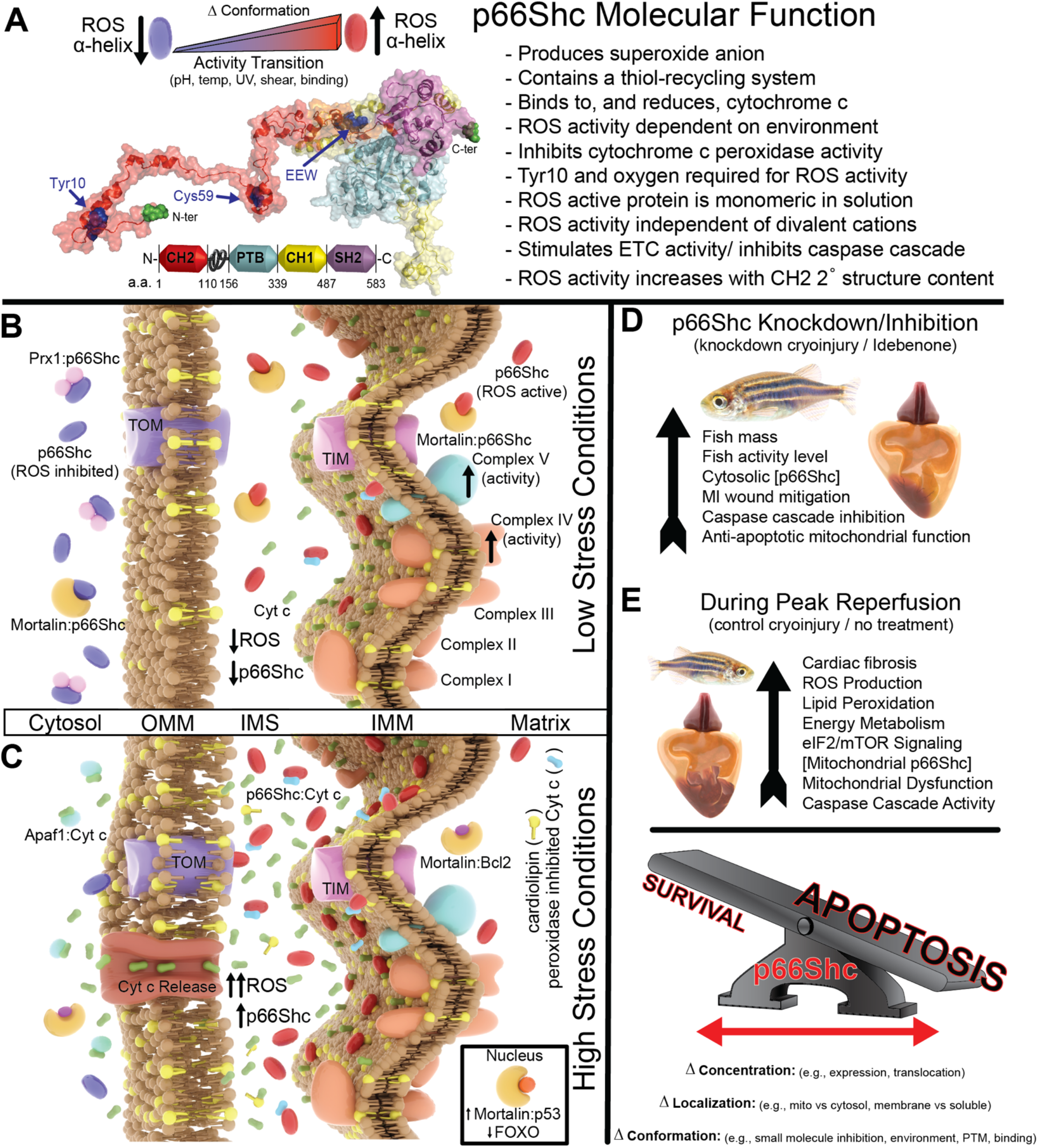
p66Shc acts as a molecular rheostat to govern apoptosis activity in a cell stress-dependent manner. (A) Summary of key p66Shc molecular findings. (B and C) Model of p66Shc function in low and high stress. Cytosolic p66Shc is mostly inactive via Prx1or mortalin binding and a stabilized inactive conformation. These effects prevent pro-apoptotic signaling. In low stress, IMS [p66Shc] is low and promotes homeostasis via basal O_2_^-^ production, enhanced ETC activity, and lipid peroxidation inhibition by stabilizing cyt c. As cell stress increases (*e*.*g*., MI), p66shc-mortalin dissociation, p53-mediated upregulation of p66Shc, and mitochondrial transport increases. Within the IMS, p66Shc undergoes ROS-activating conformational changes, increasing [O_2_^-^] until a pathological threshold is met. This causes cyt c peroxidase formation that depletes the IMM of CL and disrupts ETC activity. Unable to further pass *e*^-^s to a dysfunctional ETC, inhibit caspase activation, or prevent cyt c peroxidase formation, p66Shc activity becomes strictly pro-apoptotic. p66Shc also inhibits Foxo3A antioxidant transcription, exacerbating antioxidant defenses. In addition, apoptosis during reperfusion increases extracellular p66Shc activity by creating an acidic, lipid-rich, and warm environment with high sheer that can propagate oxidative stress to nearby cells. (D and E) Results summary of targeted p66Shc ROS inhibition via KD or small molecules (D) vs control fish (E). Collectively, data suggest inhibiting p66Shc’s O_2_^-^ production allows p66Shc’s balance between pro-apoptotic and anti-apoptotic function to favor increased myocardial survival.

### Limitations of the Study

Defining p66Shc’s activity is inherently complex as it has multiple functions, multiple isoforms, and dynamic cellular localization that drive activity. Yet, p66Shc’s diversity supports its role as a molecular “rheostat”. To better define the “Jekyll and Hyde” function of p66Shc and unite conflicting results in prior studies, we performed mechanistic studies and inhibited p66Shc ROS activity in a zebrafish MI model. Our studies demonstrate p66Shc is a multi-functional protein, like cyt c, but complete p66Shc functional characterization requires answers to the following questions: how do electrons flow through p66Shc, how does p66Shc modulate ETC activity, and how does p66Shc structure drive function?

We identified cyt c as a novel downstream *e*^*-*^ acceptor but upstream donor(s) remain unknown. Ide is a synthetic ubiquinone mimetic that we demonstrate binds and inhibits p66Shc, improving zebrafish IRI outcomes. Thus, the more hydrophobic ubiquinone may also bind p66Shc and serve as a biological electron donor.

Data presented here, and inferred by other studies, support cyt c recycling to stimulate Complex IV activity. However, it is unclear if p66Shc bypasses Complex III via cyt c reduction or if p66Shc directly interacts with Complex IV, as both paths are consistent with increased Complex IV and V activity. In addition, mito O_2_ consumption studies and membrane potential analysis could provide detailed mechanistic insights into how p66Shc impacts metabolism in response to cell stress and disease.

Since CH2 appears intrinsically disordered but responds to environmental changes by increasing secondary structure and ROS activity, we propose the CH2 domain is an environmental sensor that modulates enzyme function. The inter-domain cysteine network impacts global p66Shc conformation but specific network states’ correlation to ROS activity and whether *e*^*-*^ transfer occurs directly via Cys residues have not been defined. Also, p66Shc binds CL to stimulate ROS production, similar to cyt c, but this interaction’s broader impact requires further study. Thus, comprehensive dynamics and conformational studies may provide mechanistic insights into CH2-dependent ROS activity modulation.

## ACKNOWLEDGEMENTS

We thank Drs. Gillian Air, Paul Weigel, Jialing Lin and Jian-Xing Ma at OUHSC and John Baynes at the University of South Carolina for suggestions during these studies. This work was supported by the National Institutes of Health under the following awards: National Institute of General Medical Sciences: R01GM118599 (F.A.H.), National Heart Lung and Blood Institute: F30HL149279 (L.H.), Pilot Project funding from an Institutional Development Award (IDeA): P20GM103650, and IDeA National Resource for Quantitative Proteomics: (R24GM137786). Additional support was provided by the Presbyterian Health Foundation (F.A.H.). UltraScan development is supported by R01GM120600 (B.D.) and UltraScan supercomputer calculations were supported through NSF/XSEDE grant TG-MCB070039N (B.D), and University of Texas grant TG457201 (B.D). We acknowledge the Oklahoma Medical Research Foundation for support of their EPR facility. We thank Drs. Javier Seravalli for assistance with ICP-MS studies and Jordan C. Evans and Yuko Tsutsui for assistance with experiments, and Megan Malone-Perez for assistance in the OUHSC Fish Facility.

## AUTHOR CONTRIBUTIONS

Conceptualization, L.H., D.M.B. and F.A.H.; Methodology, L.H., J.M.H., H.S., S.M., V.S., K.M.H., J.K.F., B.D., S.D.B., D.M.B., R.T., K.D.D., F.A.H.; Software, B.D. and F.A.H.; Validation, L.H., J.M.H., S.M., V.S., K.M.H., J.K.F., S.D.B., B.D. and F.A.H.; Formal Analysis, L.H., S.M., V.S., J.M.H., S.D.B., B.D. and F.A.H.; Investigation, L.H., J.M.H., H.S., S.M., V.S., R.T., K.D.D. and F.A.H.; Resources, L.H., K.M.H., J.K.F. and F.A.H.; Data Curation, F.A.H.; Writing – Original Draft, L.H. and F.A.H.; Writing – Review and Editing, L.H., J.M.H., H.S., S.M., V.S., K.M.H., J.K.F., B.D., D.M.B., S.D.B., R.T., K.D.D., F.A.H.; Visualization, L.H. and F.A.H.; Supervision, F.A.H.; Project Administration, J.K.F., J.M.H., F.A.H.; Funding Acquisition, L.H. and F.A.H.

## STAR★Methods

### RESOURCE AVAILABILITY

#### Lead contact

Additional information and resource or reagent requests should be directed to, and be fulfilled by, the Lead Contact: Franklin A. Hays.

#### Materials availability

All plasmids, cell lines, and other materials are available on request from the Lead Contact.

#### Data and code availability

- Comparative proteomic data is provided in **Table S1** and mass spectrometry data provided in **Table S2**. Raw data files are available from the Lead Contact upon request.
- This paper does not report original code.
- There are no restrictions on data availability in this manuscript. Data and any information required for data analysis reported in this study are provided in Supplemental Information or available from the Lead Contact, on request.

### EXPERIMENTAL MODEL AND SUBJECT DETAILS

#### Zebrafish

*Danio rerio* fish heterozygous for an obligate p66Shc gene splice site, generated by ENU mutagenesis, were obtained from the Zebrafish International Resource Center (ZIRC), ZIRC catalog ID ZL12658.11. Zebrafish were housed in a 28.5°C temperature controlled aquatic colony with a 14 hour light and 10 hour dark circadian cycle. All experiments were performed consistent with pre-approved University of Oklahoma Health Sciences Center IACUC protocol 19-058-SAH. Obtained fish were genotyped and splice site females were outbred for 3 generations with *myl7:eGFP* homozygous fish to create heterozygotes for testing. WT fish were either *myl7:eGFP* homozygotes or non-splice site clutchmates from a single parental breeding pool. All genotyping procedures were performed with anesthetized fish, using 0.02% tricaine methanesulfonate (MS-222). Fish were genotyped via tail clip DNA extraction followed by PCR using these primers: TTG GCT TCC AAA GAG AGA CTT CA (forward), GTG AAT GTC GTT CTG GCT CG (reverse). High fidelity phusion polymerase (NEB) was used for all sequencing reactions. PCR reactions were then cleaned per Monarch PCR & DNA cleanup kit protocol (NEB). Purified amplicons were sequenced using the above primers. Cryoinjury was performed on young healthy adult fish between 6-8 months age. Gender effects were excluded by combining both male and female fish across experimental studies. Genotyped fish were randomly selected from a pooled genotyped tank and randomly placed into experimental group tanks until experimental groups had a minimum of 3 fish. Fish hearts were then assessed with a combination of stereomicroscopy, light microscope histological stains, fluorescent stains, whole-heart homogenate proteome analysis, and scanning electron microscopy. Fish mass and activity were measured before experimental conditions were applied and immediately before sacrifice. All animal experiments were conducted under approved protocol 19-058-SAHB from the OUHSC Institutional Animal Care and Use Committee and conforms with the *Guide for the Care and Use of Laboratory Animals, 8*^*th*^ *Edition* published by the National Institutes of Health.

#### Human cell lines

Human WI-38, PCS-201, and HS27 fibroblast cell lines from American Type Culture Collection (ATCC) were grown in 25 cm^2^ culture flasks at 5% CO2 and 37° C in Dulbecco’s Modified Eagle Medium and 10% FBS with, 100 mg/mL streptomycin, 100 units/mL penicillin and 100 μg/mL amphotericin. WI-38 cells were supplemented with 1 mM sodium pyruvate. After approximately 80% confluence, cell lines were treated with 0.25% Trypsin/EDTA solution and incubated for 15 minutes at room temperature. Trypsin/EDTA was neutralized with growth media. Cells were centrifuged at 100 x g at room temperature for 9 minutes. Media was removed and cells were resuspended in PBS to a final concentration of 2×10^6^ cells/mL before performing assays. Materials were purchased from Sigma Aldrich except FBS purchased from ThermoScientific.

### METHOD DETAILS

#### Chemicals and reagents

Source of chemicals and reagents are provided in the Key Resources table.

#### Zebrafish cryoinjury experiments and organ collection

Cryoinjury was performed as described previously (Gonzalez-Rosa and Mercader, 2012). In brief, fish are anesthetized by immersion with 0.032% (*w/v*) MS-222 and immobilized in a sponge holder. Under a dissecting microscope, a small incision exposes and opens pericardial sacs. Gentle abdominal pressure is applied to expose ventricles. Hearts were blotted dry with paper towels before a liquid nitrogen pre-cooled 0.3 mm diameter copper probe was placed on the exposed ventricle for 30 seconds, inducing a myocardial infarction. To avoid mechanical damage, the cryoprobe was held still for the cryoinjury’s duration. Fish were resuscitated, by pipetting water over their gills until a normal breathing pattern was established. Dry probes were pre-cooled by placing their tips in liquid N_2_ for a minimum of 1 minute prior to inducing cryoinjury and cleaned between surgeries with tank water and paper towels. Negative controls were also included. Negative controls were fish that received a sham cryoinjury where a room temperature probe was applied for 30 seconds to exposed ventricle, instead of a liquid N_2_ cooled probe, preventing cryoinjury. If applicable, treatments with 1 μM carvedilol, 1 μM idebenone, 1 μM rapamycin, or 500 μM NAC were started 3 days before cryoinjury. Fish were treated for 1 hour in a separate treatment tank (3 fish per tank) containing 1 L of system water each day. Therapeutics were administered in 1 mL volumes at 1,000X concentration, using pure ethanol as vehicle. Fish that did not receive a treatment were handled identically, but received only vehicle. After treatment, fish were placed in new tanks containing 3 L of fresh system water. Daily one-hour treatments were continued for one-week post-cryoinjury before fish were sacrificed.

Prior to harvesting organs, fish were anesthetized in 0.02% MS-222 and euthanized using ice-water. Original incisions were expanded and forceps were used to stabilize hearts by pinching the bulbous arteriosus before releasing hearts by cutting between the atrium and sinus venosus. Each fish represented a single biological replicate. Replicate amounts are indicated in each figure panel or legend. Reported statistical values are relative to WT fish that received cryoinjury but no treatment, unless otherwise noted. Confounding variables were excluded by performing surgery and recovery in identical locations and conditions for each surgery-recovery combination. Genotypes and treatments were not blinded. Sample sizes were determined based on statistical variance within experimental groups. A minimum of 3 fish were used in each group. Only fish that died during surgery or before tissue harvest were excluded from analysis, as their hearts were not collected.

#### ShcA cloning and mutagenesis

cDNA for human p66Shc was purchased through OriGene (Rockville, MD). ShcA sequences were amplified and placed into a modified pET52 expression plasmid, containing an N-terminal maltose binding protein (MBP) via restriction digest cloning. MBP was followed by a 6X-His tag, a 3C protease restriction site, and a corresponding DNA sequence: p66Shc WT, p52Shc WT, p46Shc WT and CH2 WT. Sequences were verified through conventional DNA sequencing with serial cloner 2.6.1 and FinchTV 1.4.0 software before mutants were generated, using WT sequences as templates. Mutants were generated via quickchange PCR (Agilent Technologies Inc.). Mutations were verified in the same manner as WT sequences. Cloning work and plasmid generation were performed with OmniMax™ cells. Plasmids containing verified sequences were transformed into Rosetta 2 (DE3 pLysS), Shuffle® T7, or Origami™ B(DE3) *Escherichia coli* cells via heat shock.

#### ShcA protein expression and purification

Freshly transformed *E. coli* cells or glycerol stocks maintained in antibiotics were cultured in Luria broth (LB) at 37° C until an OD of ∼0.6 and induced with 50 μM – 1 mM IPTG. After administering IPTG, growth temperature was lowered to 30° C. Cells were incubated for an additional 3 hours at 250 RPM. Cells were collected following a low-speed spin at 4000 x g for 30 minutes and resuspended in ice cold 20 mM TRIS-HCl pH 7.4, 150 mM NaCl, 4 mM fresh PMSF, and 1 mM DHALT protease inhibitor. Cell pellets with overexpressed ShcA proteins were used immediately or stored at -20° C in resuspension buffer until use. If frozen cells were used, they were thawed over ice and incubated with DNAse/RNase and stirred at 4° C for 20 minutes. Cells were lysed using either a Sonics Vibra Cell VCX500 sonicator with a 13 mm probe (630-0220) at 30% amplitude with 30 sec on/off cycles for 20 minutes or an Avestin C3 Emulsiflex with 3 complete passes at an average of 15,000 psi. Cell lysate was centrifuged at 26000 x g to remove cell debris. Protein purification was performed by loading cell lysate onto an in-house made maltodextrin resin column by gravity flow. Maltodextrin resin was prepared as previously described (Teichberg et al., 1988), using a 10% (*w/v*) maltodextrin solution, and stored in 20% (*v/v*) ethanol. Resin was washed with 250 mL of 20 mM Tris pH 7.4 RT, 5 mM EDTA, and 150 mM NaCl. A second wash was performed with 100 mL of 20 mM Tris pH 7.5 RT, 5 mM EDTA, and 100 mM NaCl. Elution was performed with 20 mM Tris pH 7.5 RT, 100 mM maltose, 5 mM EDTA, 50 mM NaCl, and 2 mM octyl-glucosyl neopentyl glycol (OGNPG, Anatrace). Eluted samples were cleaved from their tags using a MBP-tagged 3C protease at 4° C overnight while dialyzing with 20 mM Tris pH 7.4 RT, 5 mM EDTA, 20 mM NaCl. Digested samples were run on an amylose resin at 4° C, to remove protease. Flow-through was pooled and ran against a Hitrap Q column (5 mL) that had been equilibrated with 20 mM Tris pH 7.4 RT, 5 mM EDTA, 2 mM OGNPG, and 25 mM NaCl. This buffer was used for a single wash before eluting protein in a 1 M NaCl linear gradient. Fractions containing p66Shc (verified via SDS and, initially, mass spectrometry) were pooled. Pooled fractions were dialyzed against desired buffers. Dialyzed proteins were lyophilized for storage or used immediately. Concentrations were determined by absorbance readings at 280 nm with extinction coefficients determined by ExPASy protparam (https://web.expasy.org/protparam/). Protein identification was validated using MALDI-TOF mass spectrometry.

#### Hydroethidine superoxide measurements

Hydroethidine (HE) assays were performed in an ISS photon counting spectrofluorometer by measuring 2-hydroxyethidium (2-OH-E^+^) appearance rates, a fluorescent product produced when HE reacts with superoxide. Samples contained 20 mM MOPS pH 7.4 RT, 1 mg/mL sheared salmon sperm DNA (Invitrogen) and 10 μM diethylenetriaminepentaacetic acid (DTPA, metal chelator). DNA was added to increase fluorescence quantum yields, as previously reported (Zielonka et al., 2008). Reactions also contained 100 mM NaCl, 1 μM HE (experiments illustrating V_max_) or 10 uM HE (experiments with direct fluorescence reads), and 10 μM purified protein in MOPS pH 7.4. Fluorescence was monitored at 567 nm with an excitation wavelength of 480 nm. Assays were performed at 37° C and emissions were read continuously for an hour or until a linear slope could be determined, in the presence or absence of superoxide dismutase. These conditions represent base HE assay conditions, a control against which variables were tested. All HE assays were prepared in a similar manner but with changes to a specified variable (variables noted below). Measurements were repeated 2-4 times with unique protein samples. Average values were used for statistical analysis. In each instance, the 2-OH-E^+^ product was further isolated using reverse phase HPLC as previously described (Zhao et al., 2005; Zielonka *et al*., 2008), and shown in Figure3B, to quantify fluorescent product derived from O_2_^-^.

V_max_ was determined by normalizing fluorescence intensity across all samples to their 37° C WT counterpart (*e*.*g*., p66Shc and p66Shc mutants to p66Shc WT tested at 37° C, CH2 and CH2 mutants to CH2 WT tested at 37° C). The percent remaining reaction was determined by deriving a linear equation for each sample using initial and peak fluorescence intensity weighted by 2-OH-E^+^ yield. Activity was determined by calculating fluorescence intensity changes per minute. Normalized % remaining reaction values were plotted against activity in Graphpad Prism. V_max_ was calculated using the Michaelis-Menten non-linear regression analysis function.

Fluorescence assays testing temperature effects on ROS activity were prepared, as described above. Assays used 10 μM concentrations of WT human p66Shc or purified WT CH2 domain and were performed at 5°C intervals, starting at 17 ° C, ending at 77 °C. While determining maximal activity with respect to temperature, all other conditions were held constant. After performing temperature experiments without cardiolipin (CL), selected temperatures (22, 37, and 62° C) were repeated using either 2.5 mM or 10 mM cardiolipin. CH2 lipid effects were only tested at 37 °C with 2.5 mM CL. Detergent micelles were only tested on CH2 using 0, 1, 2, 3, 4, and 5, mM OGNPG. To test pH effects on ROS activity, protein was dialyzed (1:1,000 volume ratio) in buffers with a constant 100 mM NaCl concentration and a varying pH (buffered at room temperature) for 30 minutes before performing assays at 37 °C (2.5 and 3.0 pH buffers = 20 mM citrate-phosphate, 4.5 pH buffer = 20 mM MES, 5.5 pH buffer = 20 mM HEPES, 6.5, 7.4, and 8.0 buffers = 20 mM MOPS, 9.0 buffer = 20 mM Tris.) p66Shc and CH2 mutants were generated via site directed mutagenesis as outlined above and tested at 37 °C. Data collected on mutants used 20 mM Tris and 100 mM NaCl at pH 7.4 (RT) as buffer.

UV effects on protein activity were assessed by exposing protein to UV light ± pterin for 2 hours prior to conducting fluorescence assays at 37°C with pterin products isolated using size-exclusion chromatography. Chelator effects on CH2 were tested by performing assays at 37°C with 10 μM DTPA or a combination of 10 μM DTPA and 5 mM EDTA and comparing to samples containing no chelators. Various copper (I) and (II) concentrations were tested against CH2 ROS activity using HE assays at 37°C. Copper (I) concentrations were 0.25 to 2500 nM in 10-fold increments while copper (II) concentrations ranged from 0.25 to 2500 μM in 10-fold increments. O_2_ effects were tested by argon purging CH2 and p66Shc samples for 5 minutes prior to HE assays at 37°C or performing p66Shc assays in an anoxic chamber at 37°C. Shear effects were determined by including a stir bar at 37°C with rotation set to ∼ 1200 RPM. Thermal melt functional analysis was performed by heating p66Shc WT to 97 °C for either 5 or 30 minutes with no CL, 2.5 mM CL, or 10 mM CL. Heated protein was allowed to cool at RT for 6 hours before HE assays were conducted at 37°C.

Chemical modifications were only performed on CH2. CH2 has a single Cys and a single Tyr. Cys59 was chemically blocked by incubating CH2 with NEM (N-Ethylmaleimide, Sigma) while Tyr10 was blocked by incubating a separate CH2 batch with acetylimidazole overnight at 4°C. Final NEM and acetylimidazole concentrations were 2 mM in 1X PBS. Excess acetylating or alkylating agent was removed via extensive dialysis (3X 1:1000 volume ratio exchanges for 90 minutes each). After acetylation effects were analyzed, acetylation was reversed by incubating the same batch of acetylated protein with 10 mM hydroxylamine in 20 mM Tris-HCl with 100 mM NaCl at 4°C, overnight. Remaining hydroxylamine was removed through dialysis, as above, and tested for activity once more. Small molecule p66Shc inhibition HE assays used varying concentrations of small molecules (carvedilol, N-acetyl cysteine, idebenone, and rapamycin). Final concentrations are indicated for small molecule inhibition assays in their corresponding figures.

HE adduct ratios formed when HE reacts with H_2_O_2_ or O_2_^-^ (producing either ethidium or 2-OH-E^+^, respectively) were determined by preparing samples with 5 μM p66Shc WT, 20 mM MOPS pH 7.4, 100 mM NaCl, 10 μM DTPA, and 10 μM HE. Samples containing p66Shc were compared to samples that did not include p66Shc. Samples were incubated for 60 minutes at RT before protein was removed by passing through an Amicon Ultra centrifugal filter (30 kDa MWKO). Flow-through was immediately injected onto a C18 column for HPLC analysis. Columns were washed to elute unreacted HE, ethidium, and 2-OH-E^+^. Fluorescence emission at 567 nm was continuously monitored as previously described (Zielonka *et al*., 2008).

#### ICP-MS cation analysis

ICP-MS protein samples were analyzed at the Department of Redox Biology/Biochemistry, University of Nebraska Lincoln campus. ROS active, and purified, protein was submitted at 15 μM concentration and diluted 10-fold prior to being ran on an Agilent ICP-MS 7500cx equipped with a 96-well plate reader from Elemental Scientific Inc. (ESI. Omaha, NE). Samples were analyzed using a mixed gas mode (3.5 mL/min H_2_ and 1.5 mL/min H_3_ gas flow) to reduce polyatomic interference. Analysis settings were set to 1600 W plasma power, 7.0 mm sample depth, 0.6 L/min carrier argon flow and 0.55 L/min makeup argon flow. Samples were introduced via an injection valve at a flow rate of ∼55 uL/min, with a 50 ppb ^71^Ga internal standard. Sample dilution matrix consisted of 2% (*v/v*) nitric acid in metal-grade analytical water, which contained a 50 ppb ^71^Ga internal standard. All sample readings were estimated using an external standard mix ran under identical conditions and calculated using ChemStation software. Three unique protein samples were run with results are presented as their average.

### Electron paramagnetic resonance

Electron paramagnetic resonance (EPR) experiments were performed as previously described (Duling, 1994; Matsuzaki et al., 2011). In brief, EPR spectra were collected on a Bruker EMX spectrometer (Billerica, MA) with an X-band (∼9.78 GHz) and a 100 kHz modulation frequency, using an ER 41225SHQ high-sensitivity cavity. Parameter settings were 6.325 mW microwave power, 1.5 G modulation amplitude, 150 G scan range, 655 msec time constant. Experiments were performed by incubating 6.11 μM protein in a MOPS pH 7.4 RT buffer containing 25 mM 5-(2,2-dimehtyl-1,3-propoxycyclophosphoryl)-5-methyl-1-pyrroline N-oxide (CYPMPO), 1 mM EDTA, 50 μM DTPA, and 25 mM KH_2_PO_4_ with or without 40 U/mL of CuZn-superoxide dismutase (SOD). Reaction solutions were incubated in microcentrifuge tubes, then transferred to a quartz flat cell immediately prior to EPR evaluation. Incubation times were typically 150 minutes. Experiments were performed at RT. Spectral simulations were completed with WinSim 2002 software from NIEHS public EPR software packages (http://epr.niehs.nih.gov). Experiments were performed with protein samples from multiple preparations.

### Human p66Shc and CH2 domain analytical ultracentrifugation

p66Shc and CH2 WT and mutant oligomerization states in free solution were analyzed using a Beckman Optima XL-I analytical ultracentrifuge (AUC) at the University of Texas Health Science Center, San Antonio as described previously (Tsutsui et al., 2015). Briefly, solutions with 8.7 μM protein in 20 mM TRIS pH 7.4 RT and 100 mM NaCl were subjected to sedimentation velocity experiments at 42,000 RPM (CH2/CH2CB) or 35,000 RPM (p66Shc and p66Shc-C59S) and 20 °C. Experiments were scanned at 280 nm for p66Shc/p66Shc-C59S or 220 nm for CH2/CH2CB in intensity mode inside a 1.2 cm epon 2-channel centerpiece (Beckman Coulter). Partial specific volumes were estimated from protein sequence with UltraScan and found to be 0.7274 ml/g for p66Shc and 0.7258 ml/g for CH2CB. Data were analyzed with UltraScan-III version 4.0 release 6209 (Demeler and Gorbet, 2016). Data refinement was performed as described (Demeler, 2010), using two-dimensional spectrum (Brookes et al., 2010) and Monte Carlo analysis (Demeler and Brookes, 2008). The enhanced van Holde – Weischet analysis (Demeler and van Holde, 2004) was used to determine diffusion-corrected integral sedimentation coefficient distributions. AUC measurements were performed at the Center for Analytical Ultracentrifugation of Macromolecular Assemblies at the University of Texas Health Science Center at San Antonio and calculations were performed on Texas Advanced Computing Center’s Lonestar Cluster, University of Texas.

### Microscale thermophoresis (MST) experiments

MST experiments were carried out using standard MST capillaries in a monolith NT 115 instrument (NanoTemper) equipped with a red filter. Prior to conducting experiments, protein was labeled with 500 μM Alexa 647, per the manufacturer’s instructions. Experiments took place in 20 mM MOPS and 100 mM NaCl, pH 7.4. For p66Shc binding studies, a 24 nM concentration of labeled protein was incubated with 16 tubes of 2X serially diluted ligand. Starting concentrations of each ligand for p66Shc studies were 15.6 μM carvedilol, 31.3 μM Cu^1+^, 500 μM Cu^2+^, 1 mM ferric cyt c, 900 μM ferrous cyt c, 1 mM NAC, 0.5 μM rapamycin, 0.75 μM Idebenone, and 0.5 mM iso-cyt c (constitutively active cyt c peroxidase). Studies aimed at determining p66Shc binding specificity for carvedilol, idebenone, N-acetyl cysteine, and rapamycin were done using CH2 (only present in p66Shc isoform). For small molecule specificity, CH2 was labeled with Alexa 647 and experiments used 34 nM CH2. Ligands were prepared in 16 tubes, using a 2X serial dilution. Initial concentrations were as follows for CH2 binding analysis: 31.3 μM carvedilol, 2 μM Idebenone, 500 μM NAC, 250 μM rapamycin. For cyt c specificity, labeled CH2 concentration was 75 nM and all cyt c forms (reduced, oxidized, and peroxidase active) had 500 μM starting concentrations. Lastly, 75 nM CH2 was also tested against Cu^1+^ and Cu^2+^. Starting copper concentrations were 0.5 mM for each copper species. All measurements have an N value ≥ 3 with final K_D_ values reported as their averages. K_D_ values were determined using non-linear regression in Graphpad Prism software. All binding experiments were performed in a 25 °C temperature-controlled environment. Mortalin-BCL-2 interactions were analyzed in 20 mM MOPS pH 7.4, 10% glycerol, 100 mM NaCl at 37°C. For these experiments mortalin was labeled with Alexa 647 dye (NanoTemper) and conducted with 50 nM mortalin. p66Shc-mortalin interactions were performed by labeling mortalin with NT-647 prior to running assays with 25 nM of fluorescently labeled mortalin in 20 mM MOPS pH 7.4, 1.5mM OGNG, 100 mM NaCl at 25 °C. For MST experiments, MST power was set to 40% with excitation power adjusted to optimize signal yield.

### Interaction of p66Shc with cytochrome c analyzed by surface plasmon resonance (SPR)

SPR analysis of ferric cyt c and human p66Shc binding was conducted using a SensiQ Pioneer at a controlled analysis temperature of 25 °C in 20 mM MOPS pH 7.4, 100mM NaCl, 1 mM EDTA, 50 μM DTPA, 1mM octyl glucose neopentyl glycol (OGNG) (Buffer B). COOH2 planar dextran sensor chip (SensiQ Technologies Inc.) was twice pre-conditioned in 10 mM HCl, 50 mM NaOH, and 0.1 % *w/v* sodium dodecyl sulfate injected over 30 sec. Sensor surface was activated by 4 min-injections of 20 mM EDC/5 mM NHS over flow channels (FC) 1-2. Ferric cyt c in 10 mM sodium acetate at pH 6.5 was immobilized on FC1 via amine coupling using 5 min injection time. FC1-2 was deactivated via 4 min injection of 1 M ethanolamine at pH 8.0. Human p66Shc binding assays were performed using OneStep™ injections in duplicate of 1 μM p66shc in Buffer B. Data analysis was performed using QDat software. All data were double referenced to the FC2 reference channel as well as three buffer blank injections. The obtained data were fitted to 1:1 Langmuir binding model.

### *In silico* p66Shc modelling and molecular dynamics simulations

The isolated PTB domain structure (residue 111-317 in p66Shc numbering) was solvated with TIP3P water molecules using the SOLVATE module in VMD software package (Humphrey et al., 1996). NaCl (100 mM) was added to maintain neutrality of the system using the AUTOIONIZE plugin. Mutations to construct p46Shc or E132Q/E133Q double mutation were carried out using the MUTATOR plugin. The final dimension of the system was 132.2 Å x 92.9 Å x 74.7 Å. All simulations were performed using NAMD (Phillips et al., 2005) combined with CHARMM36 force field (Vanommeslaeghe et al., 2010)as an NPT ensemble with 2 fs time step. Energy minimization of the system was carried out with 10000 steps. Constant pressure (1 atm) was maintained using the Nosé-Hoover Langevin piston with 100 fs piston period and 50 fs damping time constant. Langevin dynamics with a damping coefficient of 1 ps^-1^ was used to maintain temperature at 298 K. Non-bonded interactions were cut off after 12 Å with application of a smoothing function after 10 Å. The particle mesh Ewald method was used to calculate electrostatic interactions with grid spacing of 1.0 Å, and the periodic boundary condition was used. Figures were prepared using Chimera (Pettersen et al., 2004). The full-length molecular model presented in **Figure 3A** and **Figure 5B** was generated using version v1.0.0 of RoseTTAFold as described previously (Baek et al., 2021).

#### Cytochrome c redox assays

Ferric cyt c was prepared by dissolving horse heart cyt c (Sigma) in 20 mM MOPS 100 mM NaCl and frozen in liquid N_2_ before storing at -80 °C until needed. Ferrous cyt c was prepared by adding 1 mM sodium ascorbate to dissolved ferric cyt c. Sodium ascorbate was removed via overnight dialysis (≥ 100X) at 4 °C and prepared for storage with liquid N_2_. Cyt c reduction values were determined by evaluating cyt c absorption ratios for 550 nm and 521 nm wavelengths, as described (Liu et al., 2014). A_550_/A_521_ absorbance ratios corresponding to fully reduced or oxidized cyt were set to 1.8 or 0.795, respectively. Reported values were calculated using linear regression of reduced to oxidized ratios. Samples were blanked immediately after adding ShcA protein to a solution containing cyt c. Absorbance measurements were taken every 30 minutes from 380 to 650 nm. Reactions took place in a 20 mM MOPS and 100 mM NaCl solution with a pH of 7.4. Final concentrations were 10 μM ShcA protein and 60 μM ferric, or ferrous, cyt c. Measurements were performed 3 times and statistical analysis used their average values. For cyt c studies where absorption spectra are shown, 20 uM cyt c concentrations were used. Cyt c reduction assays performed with small molecules were conducted in a like manner but with final concentrations of either 25 nM carvedilol, 50 μM NAC, 25 nM idebenone, or 25 nM rapamycin.

#### Cytochrome c peroxidase functional assays

20 μM ferrous cyt c was incubated for 30 minutes at RT in a 20 mM MOPS buffer containing 100 mM NaCl and 100 μM cardiolipin, pH 7.4. Cyt c peroxidase activity was measured at RT using 10 mM guaiacol and various concentrations of H_2_O_2_ by measuring tetra-guaiacol formation at 470 nm (E^mM^ = 26.6). Measurements were started immediately after H_2_O_2_ addition. Inhibition assays were performed by using different concentrations of p66Shc in peroxidase assays and monitoring tetra-guaiacol accumulation. Cyt c peroxidase assays were also performed using amplex red with CH2. Amplex red is an H_2_O_2_-sensitive dye, commonly used to assess H_2_O_2_ levels (Deshwal et al., 2018). CH2 solutions were incubated with iso-cyt c for 1 hour before adding amplex red and H_2_O_2_. Samples were blanked immediately after adding H_2_O_2_ before spectrophotometrically measuring resorufin formation at 572 nm. Final concentrations for these assays were: 10 μM CH2, 1 mM amplex red, 100 μM iso-cyt c, 20 mM MOPS, 100 mM NaCl and 500 μM H_2_O_2_. Experiments were conducted a minimum of 3 times and statistical analysis was performed using their averages.

#### Circular dichroism

2.5 μM ShcA protein was dialyzed into 20 mM Tris pH 7.4 RT and 100 mM NaCl with no CL, 2.5 mM CL, or 10 mM CL. Solutions were analyzed in a JASCO J-715 spectropolarimeter with a 348WI temperature controller. Protein was sealed in a 0.1 cm path length cuvette. Proteins were scanned 3 times from 260 to 190 nm at a rate of 10 nm/min. Scans started at 2 °C and ended at 97 °C by 5 °C increments. A final set of scans was performed at 37 °C, post-thermal melt to determine ShcA refolding characteristics under slow cooling conditions (overnight incubation at RT). Protein and blank samples were incubated at each temperature for at least 2 minutes before performing a scan. All scans were performed 3 times. Reported values are average sample signal minus average corresponding buffer signal. Secondary structure composition was determined by Contin structural analysis program of the fourth reference data set via DichroWeb (http://dichroweb.cryst.bbk.ac.uk).

#### Mass spectrometry, cysteine interactions

Methods were adapted and modified from a previously published protocol (Sechi and Chait, 1998). p66Shc WT at 10 μM concentration was incubated at RT for one hour in a 20 mM MOPS 100 mM NaCl pH 7.4 buffer with either 10 mM dimedone (Sigma) or NEM (Pierce) (**Table S1**). Flash freezing with liquid N_2_ prevented further reactions and was stored at -80 °C until sample was sent for MS analysis. Disulfide bonds were identified by reducing p66Shc with 5 mM DTT at 37 °C for 1 hr. prior to incubating with 15 mM iodoacetamide (Sigma), without light exposure, for 30 minutes at RT. Thus, initially free cysteines were alkylated by NEM and disulfide bonded cysteines were labeled with iodoacetamide. Alkylated p66Shc was transferred to a 50 mM ammonium bicarbonate solution via buffer exchange and was acidified prior to overnight pepsin (Promega) digestion at 37 °C using a p66Shc:pepsin ratio of 1:50, w/w. Dimedone labeled samples were also buffer exchanged into 50 mM ammonium bicarbonate prior to overnight digestion at 37 °C with trypsin (Thermo-Scientific) at a p66Shc:trypsin ratio of 1:50, w/w. LC-MS^E^ analysis to generate peptide maps were done on a Waters Acquity UPLC H-Class in line mass spectrometer with a Waters Synapt G2-S HDMS™. The spectrometer was calibrated weekly with [Glu1]-fibrinopeptide B (GFB) and green fluorescent protein (GFP) was used in the lockspray channel. 50 ng of each sample were injected onto Waters Acquity UPLC HSS T3 with a 1.8 μm particle size on a 75 μm by 150 mm column held at 65 °C. Peptide fragments were eluted with a steady 0.1% formic acid (Sigma) and 5-40% acetonitrile (Fisher Scientific) gradient for 54 minutes. Mass spectrometer resolution mode was set to positive ion and had a detection range of 50 to 2000 m/z. Remaining parameters follow: capillary voltage = 3.0 kV, sampling cone voltage = 35.0 V, source temperature = 80 °C, desolvation temperature = 150 °C, cone gas flow = 500 L/h, low collision energy = 4.0 eV, elevated energy ram =20-50 eV. Generated data were processed with Biopharmalynx version 1.3 software. During disulfide bond identification, methionine oxidation and cysteine carbamidomethylation were set to variable modifications. Searches were conducted for scrambled disulfides, with two disulfide bonds maximum per peptide. Dimedone analysis used methionine oxidation and cysteine-dimedone adducts (MW = 138.0681 Da) as variable modifications. Mass tolerances were set below 30 ppm. Identified peptide fragments were confirmed by MS^E^ spectra with a minimum of five matched b/y fragment ions.

#### Mass spectroscopy, proteomics

Resected heart samples were homogenized in RIPA buffer (Thermo) using a pestle microtube homogenizer (Fisher Scientific) for 2.5 minutes. Samples were stored at -80°C until sent to IDeA National Resource for Quantitative Proteomics, for sample processing and data collection. Samples were reduced, alkylated, and purified by chloroform/methanol extraction and followed by sequence grade porcine trypsin (Promega) digestion. Protein fragments were separated using reverse phase XSelect CSH C18 2.5 um resin (Waters) on an in-line 150 × 0.075 mm column with an UltiMate 3000 RSLCnano system (Thermo). Elution was performed with a 90 min gradient from 98:2 to 65:35 buffer ratio (0.1% formic acid/0.5% acetonitrile: 0.1% formic acid, 99.9% acetonitrile, respectively). Fragments were then ionized via 2.2kV electrospray and subjected to mass spectrometric analysis on an Orbitrap Fusion Lumos mass spectrometer (Thermo). Data was acquired in FTMS analyzer profile mode with a resolution of 120,000 and a range of 375 to 1500 m/z. Following HCD activation, MS/MS data were collected with an ion trap analyzer in centroid mode and normal mass range. Precursor mass-dependent normalized collision energy was between 28.0 and 31.0. Proteins were identified against UniprotKB *Dana rerio* database (Oct 2020) using MaxQuant (Max Planck Institute, version 1.6.17.0). Label free quantification had a parent ion tolerance of 3 ppm and a fragment ion tolerance of 0.5 Da. Scaffold Q+S (Proteome Software) verified MS/MS based peptide and protein identifications. Identifications were only accepted if they could be established with less than 1.0% false discovery and contained at least 2 identified peptides. Protein probabilities were assigned by the Protein Prophet algorithm (Nesvizhskii et al., 2003). Resultant proteins were ranked according to fold-change (**Table S3**) with follow-on functional, and disease, analysis performed using Ingenuity Pathway Analysis (IPA, Qiagen). IPA uses a Fisher’s combined probability test to determine p-values for each network and significance of linkage between observed pathways. Comparative analysis was performed across numerous samples for control (sham) ± treatment, knockdown ± treatment, and wildtype ± treatment as noted in the raw data file (**Table S3**).

#### Whole heart light microscopy injury analysis

Immediately after heart resection, hearts were rinsed with 1X PBS and images were taken under a dissecting microscope for light microscopy injury measurements using ImageJ software (1.8.0). A minimum of 3 fish were used in each treatment group.

#### Scanning electron microscopy

Hearts for each treatment group were placed in 10 mL of ice-cooled 1X PBS (pH 7.4) fortified with EM-grade glutaraldehyde (4% *w/v*). Samples were fixed overnight on a rocker in a 4°C cold room. Samples were washed 3 times in 1X PBS for 10 minutes to remove excess fixative prior to post-fixation. Post-fixation used 1 mL of 1% OsO_4_ in PBS via heart immersion. Hearts were then washed 3 times in de-ionized water before sequential alcohol dehydration with 25%, 50%, 70%, 85%, 95%, and 100% (3X) ethanol solutions. Dehydrated samples were critical point dried before mounting to SEM sample holders. Once mounted, hearts received a 5 nm AuPd sputter coat before imaging on a Zeiss Neon dual-beam HR-scanning electron microscope. Settings for individual images are indicated on each image. SEM images were analyzed with image j software (1.8.0) to determine active injury areas, total injury areas, and total heart areas. Cell classifications were based on characteristic differences in morphology that correspond to cell type and activity status. All analyses included 2-5 technical replicates per heart with a minimum of three hearts analyzed per parameter. Averaged technical replicates were used to represent a single biological replicate. Reported values are from a minimum of 3 averaged biological replicates.

#### Weight ratios and physical activity

Fish were recorded in their tanks for a minimum of 60 seconds and were weighed after measuring their activity but prior to inducing cryoinjury. Immediately before sacrificing fish, their activity and weights were recorded once more. Physical activity is presented as average distances a treatment group traveled in 60 seconds. Weight is expressed as the average ratio of pre-surgical to post-surgical weight for a given treatment group. All treatment groups had a minimum of 3 replicates.

#### Heart preparation for histology

Harvested hearts were grouped by treatment and immediately placed in 10 mL of 10% neutral buffered formalin (VWR). Samples were fixed overnight on a rocker in a 4°C cold room. Hearts were rinsed with 1X PBS, pH 7.4, for 10 minutes before embedding in Tissue-Plus OCT (Fisher Scientific) or paraffin. Hearts were cut in 6 μm serial sections (Thermo Scientific CryoStar NX70 and stained with MitoSox (Thermo), H&E (Thermo), or Trichrome (Polysciences). Differences in histology were quantified using ImageJ software. Slide images were collected by an Olympus VS120 virtual slide system for H&E and trichrome slides. MitoSox slide images were taken on a Nikon Eclipse Ts2 fluorescent microscope. Technical replicates (3-6 per parameter) were averaged to represent effects on a single heart. A minimum of 3 hearts were used for all measured parameters

#### Histology, superoxide

Superoxide staining was performed using 5 μM MitoSox, as previously performed in fixed cardiac tissue (Valls-Lacalle et al., 2016). MitoSox is a HE-derived dye that targets mitochondria and is selective for superoxide (Deshwal *et al*., 2018; Kaludercic et al., 2014; Zhao *et al*., 2005). Staining was performed in darkness, for 20 minutes, at RT. Staining was followed by one minute washes (5X) in 1X PBS before mounting slides via ProLong™ Gold antifade mountant with DAPI counterstain. Slides were concealed from light and set overnight, on a level surface and immediately imaged the following day. Fluorescence intensities were measured with NIS-Elements BR software.

#### Histology, H&E

H&E staining was performed with a Harris’ hematoxylin and eosin protocol. Briefly, slides were deparaffinized with 3 xylene washes (3 mins) followed by an alcohol hydration gradient of 100% (2X), 95% (2X), 80%, and 70% ethanol (15 dips each) before washing in deionized water and staining in Harris hematoxylin (6 mins). Samples were then rinsed in deionized water (5X), differentiated in 1% acidic alcohol (2 dips), rinsed in deionized water (6X) and blued by 0.2% ammonia water. Samples were then rinsed again in deionized water (2X), stained using eosin-y (30 secs), rinsed in 95% (2X) and 100% (3X) ethanol (10 dips each). Slides were cleared in 3 xylene changes and a single xylene substitute change (20 dips each) before mounting with xylene based medium.

#### Histology, trichrome

Trichrome staining was performed per the manufacturer’s instructions (Polysciences) by incubating slides in preheated Bouin’s fixative for 1 hour. Fixative was removed with running water (5 mins) before staining with Weigert’s hematoxylin (10 mins). Hematoxylin was removed via running water (5 mins) before staining samples with Biebrich Scarlet – acid fuchsin (5 mins). Slides were prepared for 10-minute incubation in phosphotungstic/phosphomolybdic acid by rinsing in deionized water (3 changes). After incubating with phosphotungstic/phosphomolybdic acid, slides were rinsed in deionized water for another 3 changes and transferred to aniline blue for 5 mins. Slides were rinsed in deionized water before transferring to 1% acetic acid (1 min). Slides were dehydrated and mounted as described above for H&E staining.

#### Caspase cascade inhibition

After cells were resuspended in PBS to a final concentration of 2×10^6^ cells/mL, 20,000 cells were added to each well of a 96-well solid white, flat bottom plate (Costar). Caspase-Glo 3/7 and Caspase-Glo® 9 assays (Promega) measured caspase activity. Experiments were conducted according to the manufacturer’s protocol. Caspase-Glo® 3/7 buffer and lyophilized substrate were brought to RT before they were thoroughly mixed. Substrate solution was mixed in a 1:1 ratio with 25 μL of cells. After 60 minutes, plates were read on a FLUOstar Omega Filter-based multi-mode microplate reader. Bioluminescence was used in place of fluorescence due to quenching effects of BSA on fluorescent-molecule cleavage reactions. For analysis using purified caspase 3, purified human caspase 3 (Enzo Life Sciences) was added to a 96-well solid white, flat bottom plate (Corning) at 0.05 U or 0.1 U per well. After caspase was added, plates were sealed to prevent evaporation and incubated at RT for 60 minutes. 25 μl of Caspase-Glo® 3/7 assay reagent (Promega) were added to 25 μl of cells. After 60 minutes, plates were read on a FLUOstar Omega Filter-based multi-mode microplate reader. Reported results are derived from 5 independent experiments with each experiment having 3 technical replicates per measured value.

#### Complex IV/V activity in the presence of purified p66Shc

Enzymatic activity of oxidative phosphorylation complexes were measured, according to manufacturer’s instructions, using MitoCheck Complex Activity Kits (Cayman Chemical, Ann Arbor, MI) 700990 for Complex IV and 701000 for Complex V. Mitochondria were isolated from mouse heart tissue as previously described with modifications as noted below (Vadvalkar et al., 2013; Vadvalkar et al., 2017) and used immediately. Briefly, heart tissue was perfused with ice-cold extraction buffer (EB, 5.0 mM MOPS pH 7.4, 210 mM mannitol, 70 mM sucrose, and 1.0 mM EDTA) via injection into the left ventricle. Following resection, hearts were suspended in EB and minced into small pieces with sharp scissors. Tissue was allowed to settle, blood decanted, and immediately rinsed with EB. Minced heart was then mixed with fresh EB at a ratio of ∼1 g heart weight / 5 ml buffer. Homogenate was prepared following 5 passes through a Potter-Elvehjem tissue grinder. Homogenate was centrifuged at 600 x g for 5 minutes to pellet large debris followed by passing the supernatant through 300 μm mesh into a fresh, ice-cold, centrifuge tube. Pass-through sample was then spun at 10,000 x g for 10 min, mitochondrial pellet washed in 8 ml of EB, and centrifuged a second time at 10,000 x g for 10 min. The resulting mitochondrial pellet was resuspended in 300 μl of EB. Total mitochondrial protein concentration was determined by the bicinchoninic acid assay (ThermoFisher) using BSA as a reference standard. Mitoplasts were then prepared using digitonin as previously described (Saks et al., 1985). The outer membrane was solubilized using EB containing 6 mg/ml digitonin to a final concentration of ∼25 mg mitochondrial protein / ml and mixed with continuous stirring for 15 min. Mitoplasts were collected following a 10 min spin at 8,000 x g and gently resuspended in EB to produce a final concentration of 5.0 mg / ml. Downstream coupled activity assays were completed within 4 hours of mitoplast preparation. All centrifuge steps were at 4 °C with interim samples stored on ice in pre-chilled tubes with care taken to avoid direct contact sample heating. All glassware was rinsed with 70% (*v/v*) ethanol followed by distilled water to ensure absence of detergent prior to use. For each assay, 75 μg of mitoplast was added to 10 μM of purified wildtype p66Shc, p66Shc-Y10A, or p66Shc-C59S and incubated for 1 hour using assay buffers provided in each kit. A lower, 2 μM, protein concentration was also used for Complex IV assays as indicated in **Figure 4E**. Absorbance (550 nm for Complex IV, 340 nm for Complex V) was monitored over 30 minutes with final activity measured as the change in absorbance over 1 minute (linear change). Each assay was performed multiple times (14 times for Complex IV, 8 times for Complex V) using different protein and mitoplast preparations. 1 mM KCN (Complex IV inhibitor) and 10 μg/ml oligomycin (Complex V inhibitor) were used as Complex IV and V negative controls, respectively. Data normalized to BSA-vehicle controls. To prevent Complex I interference, 1 μM rotenone was added to Complex V assays at the same time as purified protein.

### QUANTIFICATION AND STATISTICAL ANALYSIS

GraphPad Prism software calculated statistical significance. Statistical tests were unpaired two-tailed t tests, assuming Gaussian distribution during tests. No inclusion/exclusion criteria were pre-established. MST data points were excluded if MO.control software indicated capillary tube fluorescence values were inconsistent, if aggregates were detected in a capillary, or if baseline drift effects were observed. Any other excluded data points were identified via ROUT as outliers within GraphPad Prism prior to exclusion. p-values < 0.05 were considered significant. P values < 0.05, 0.01, 0.001, 0.0001 are indicated by 1, 2, 3, or 4 asterisks, respectively. Values are presented as mean ± SEM or SD, as noted in figure legends. Detailed statistical values are reported in **Table S3**.

