## Supplemental Information for "p66Shc is an apoptotic rheostat whose targeted ROS inhibition improves MI outcomes"

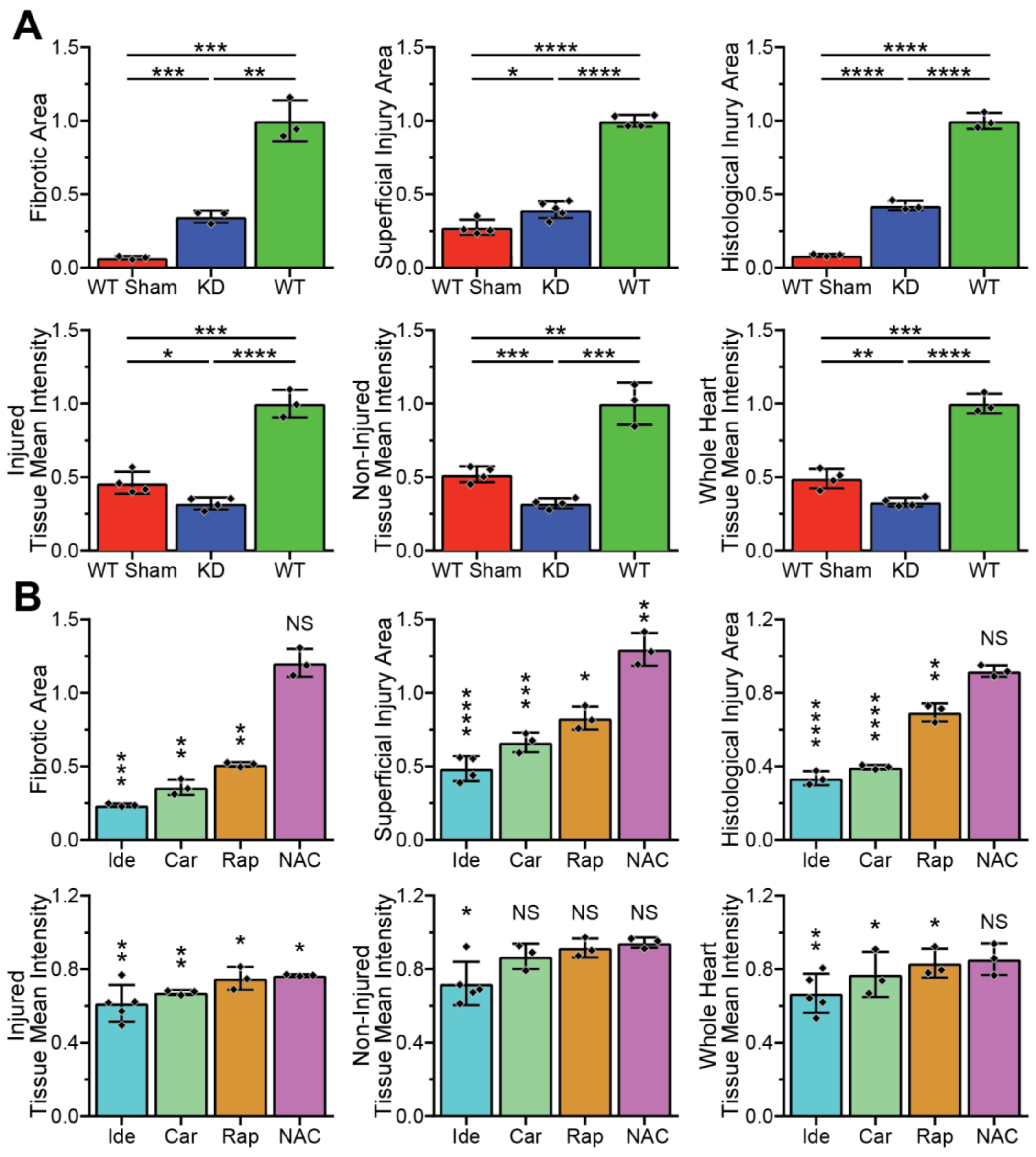


**Haslem, *et al.* - Supplemental Figure 1**

**(A-B) Trichrome (fibrotic area), stereomicroscopy (superficial injury), H&E (histological injury area), and MitoSox (mean intensity) ROS measurements for (A) KD, WT, and WT sham treatment groups or (B) WT fish treated with Ide, Car, Rap, or NAC.** Significance determined via t-tests, as indicated in (A) or against “WT” results in (B). Shown as mean ± SD. N=3-5.

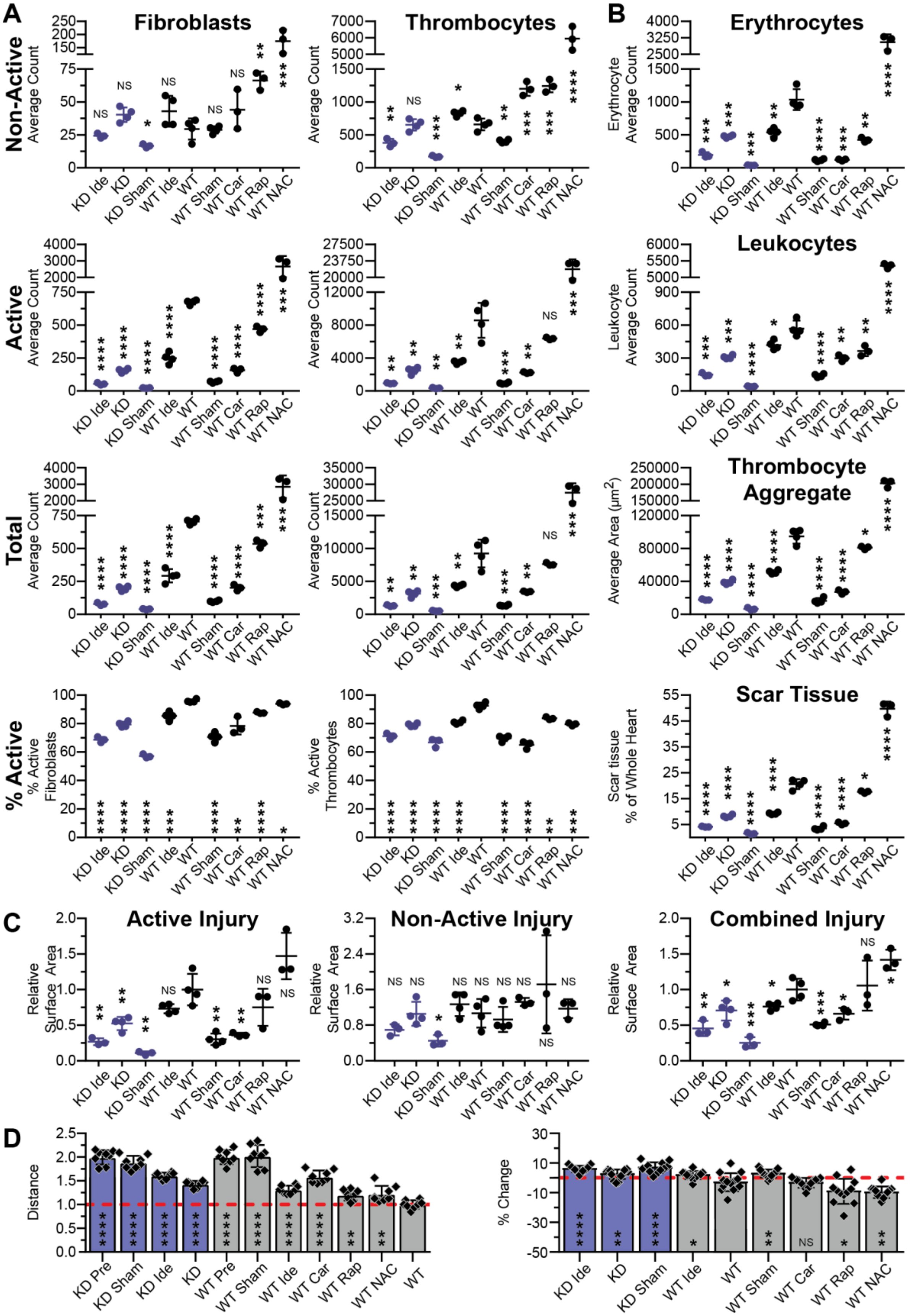


**Haslem, *et al.* - Supplemental Figure 2**

**(A-B) SEM cell counts and areas at cryoinjury site.** (A) Active, total, and % active counts for fibroblasts and thrombocytes. (B) Erythrocytes (counts), leukocytes (counts), thrombocyte aggregates (area), and scar tissue (area). Significance determined via t-tests, against “WT” results. Shown as mean ± SD. N=3-4.

**(C) Active, non-active, and combined injury areas.** Areas were measured via SEM**.** Active injury areas are closest to the initial injury site and had distinct morphological changes and high cell counts. Non-active regions had primarily morphological alterations with small differences in cell counts. Significance determined via t-tests, against “WT” results. Shown as mean ± SD. N=3-4.

**(D) PMI physical activity.** Physical activity was measured as average distance travelled per minute. Significance determined via t-tests, against “WT” results. Shown as mean ± SD. N=8.

**(E) PMI body weight ratio.** Body weight was measured immediately before surgery and heart resection. Significance determined via t-tests, against “WT” results. Shown as mean ± SD. N=10-15.

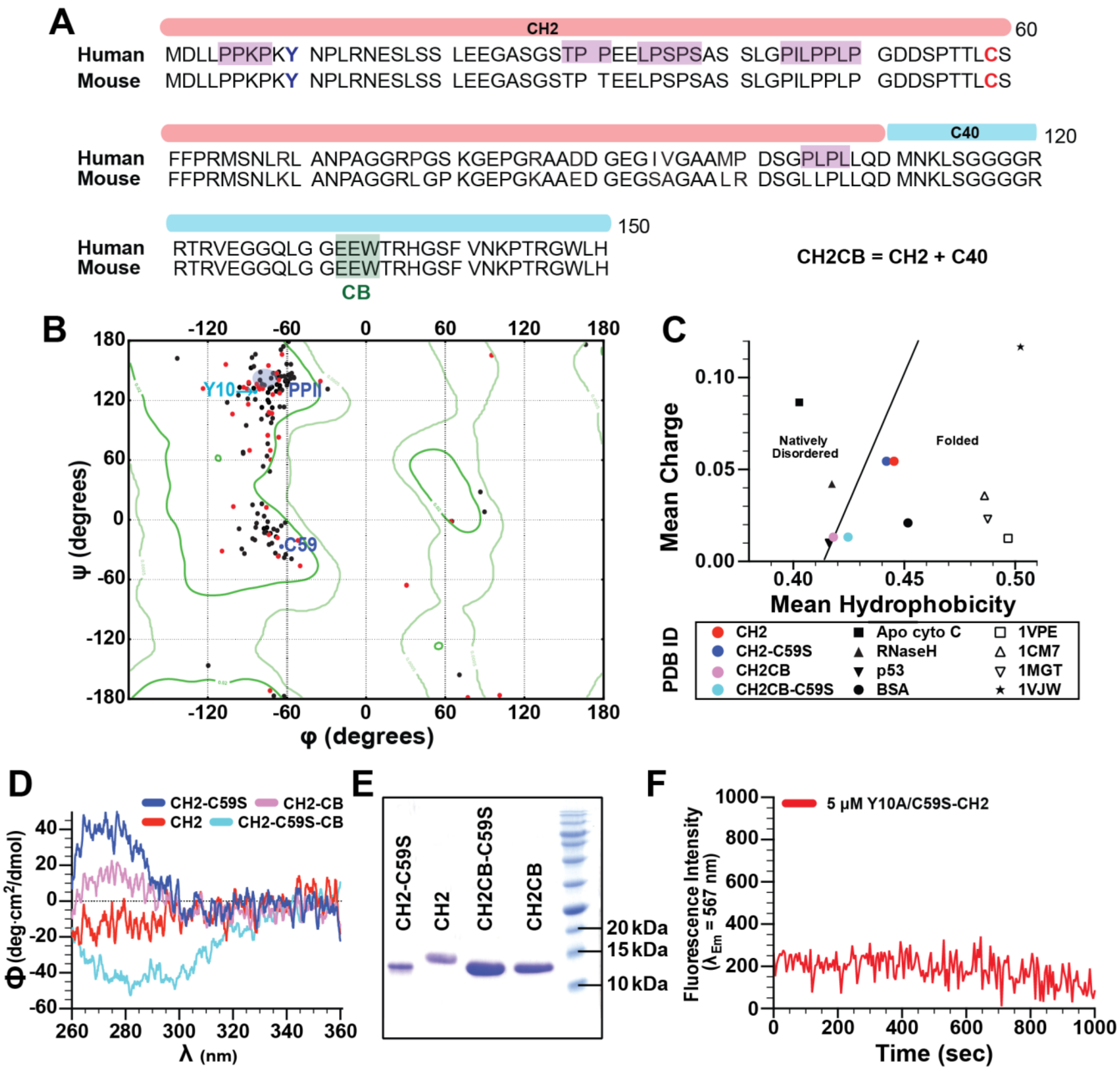


Haslem, *et al.* - Supplemental Figure 4

**(A)** The amino acid sequence alignment of N-terminal segment of human and mouse p66shc. Purple-colored boxes indicate predicted $PP_2$ helical regions by SPIDER2 (Heffernan et al., 2015). A green-colored box shows proposed cytochrome C binding region, E132, E133, and W134. Cys59 is indicated in red bold

**(B)** A Ramachandran map of human C40 region (residue 1-150). Predicted phi (φ) and psi (ψ) angles at each amino acid were obtained using SPIDER2 (Heffernan *et al.*, 2015). Black and red dots show φ and ψ angles for residues in CH2 domain (residue 1-110) or the C-terminal 40 residue extenstion (residue 111-150), respectively. Green contoured lines show favored φ and ψ angle distribution taken from 500-structure high-resolution database with B-factor <30. φ and ψ angle for Tyr10 and Cys59 are shown as cyan and blue dots, respectively (Lovell et al., 2003). Allowed φ and ψ angle for polyproline type 2 helix is shown as a shaded-purple region.

**(C)** Uversky plot for CH2 domain constructs used in this study. Mean charge and hydrophobicity of each indicated protein were obtained from a previous study and by using PONDR (Uversky et al., 2008). The boundary between folded and natively disordered region (black line) is defined in a previous study (Uversky et al., 2000). Mean charge and hydrophobicity of various thermostable proteins (indicted as PDBID number) are obtained from (Uversky et al., 2008).

**(D) 2° structure environment around aromatic residues is altered in CH2 relative to CH2CB.** Near-UV CD spectra of each indicated construct (50 μM) in 20 mM sodium phosphate, 50 mM NaCl, at pH 7.4, 25°C. CH2-C59S has a more intense CD band in near-UV CD spectrum than that of wild-type CH2CB. In the CH2 domain, aromatic residues are found at Y10, F61, and F62. In particular, Y10 is located adjacent to a predicted $PP_2$ helix region (**Figure S4A-B**) that may modulate the orientation of the Y10 side chain depending on conformational state and local environment. We hypothesize that the C59S mutation in CH2 domain primes Y10 conformation for ROS production by affecting distal $PP_2$ region spanning from residue 5 to 8.

**(E) SDS-PAGE of purified human CH2 domain constructs used in this study.** 15 % reducing SDS-PAGE of purified constructs used in this study. The expected molecular weight of CH2 and CH2CB is 11.2 kDa and 15.6 kDa, respectively. All constructs run anomalously on SDS-PAGE. (from right to left) Lane 1: molecular weight ladder, 2: CH2CB (residue 1-150), 3: CH2CB-C59S, 4: CH2 (residue 1-110), 5: CH2-C59S.

**(F) Y10A mutation abolishes ROS production in CH2-C59S.** HE fluorescence spectroscopy experiment of CH2-Y10A/C59S double mutant (5 μM) at $\lambda_{ex}$ = 480 nm, $\lambda_{em}$ = 567 nm, pH 7.4.

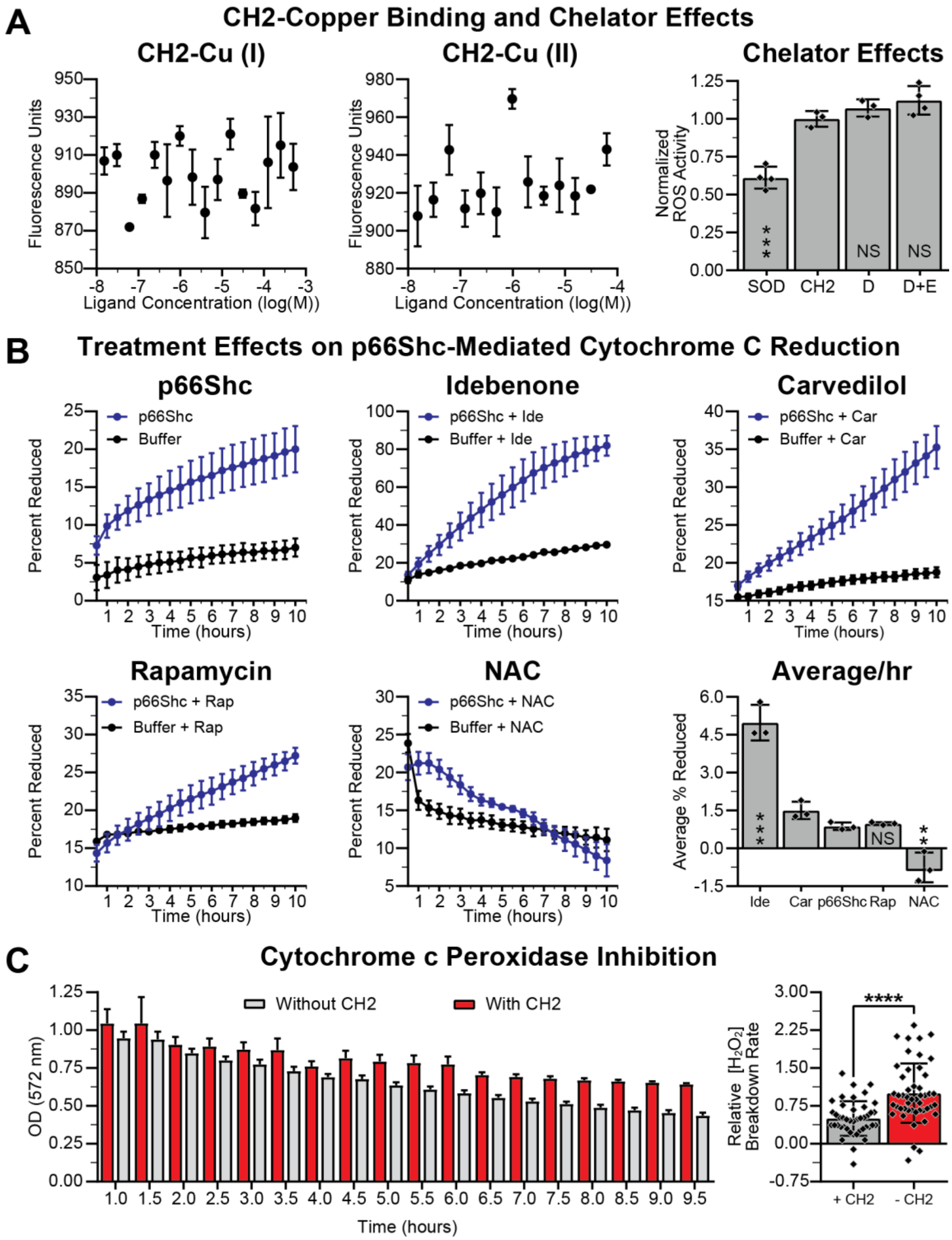


**Haslem, *et al.* - Supplemental Figure 3**

**(A) MST CH2:copper binding results and chelator effects.** No binding was observed and neither DTPA “D” nor EDTA “E” had significant effects on ROS activity. Significance determined via t-tests, against “CH2” results. Shown as mean ± SD. For binding, N=2. For ROS activity, N=3.

**(B) Treatment effects on p66Shc-mediated cytochrome c reduction.** p66Shc was incubated with oxidized cyt c and the indicated therapeutic. Significance determined via t-tests, against “p66Shc” results. Shown as mean ± SEM. N=3.

**(C) Cytochrome c peroxidase inhibition.** CH2, amplex red, peroxidase active cyt c, and $H_2O_2$ were co-incubated and OD at 572 nm was measured. Significance determined via t-tests. Shown as mean ± SEM. N=3 (left panel) and N=43 (right panel).

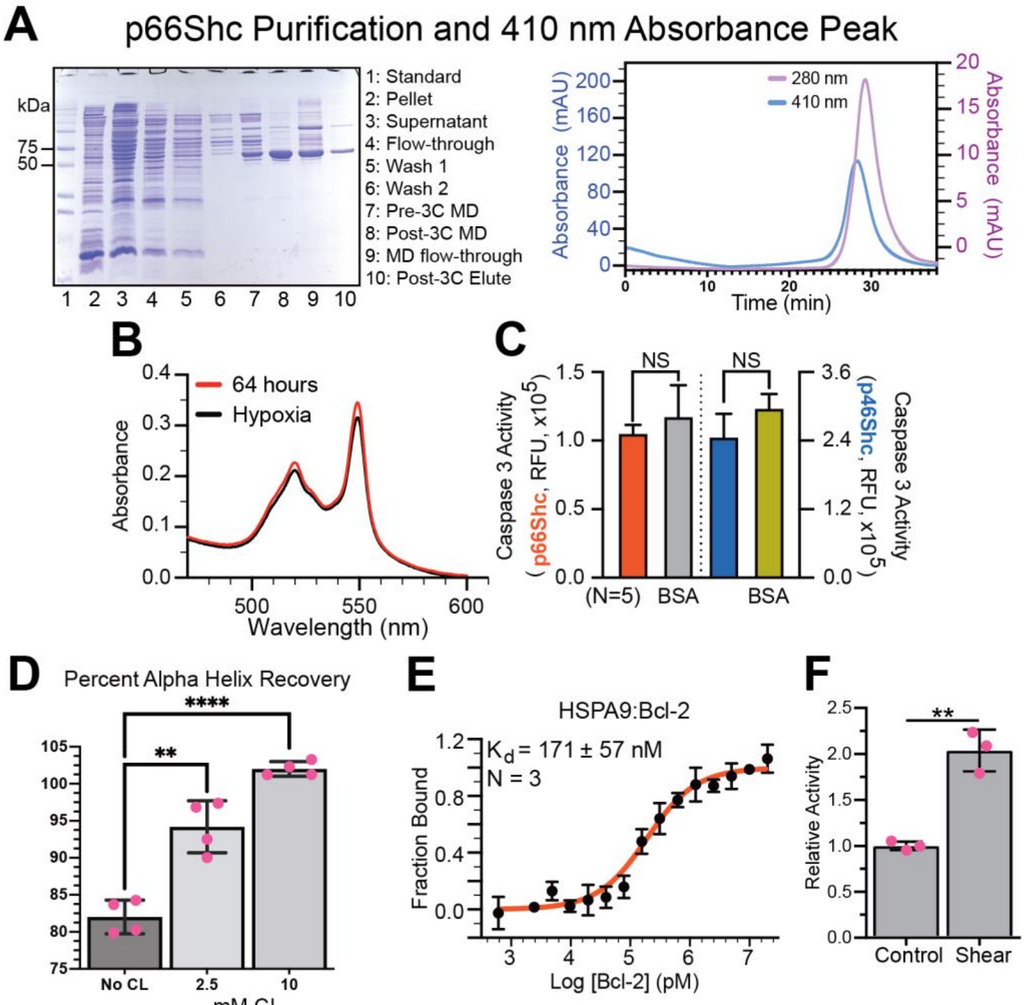


**Haslem, *et al.* - Supplemental Figure 5**

**(A) p66Shc purification.** SDS-PAGE of purified p66Shc with molecular weight marker as indicated. Purification scheme as indicated in STAR Methods with final purified p66Shc sample ("Post-3C Elute") ran on a Sephadex G-25 size exclusion column at 0.34 ml / min following absorbance at 280 nm (light blue) and 410 nm (light purple).

**(B) p66Shc reduces cyt c.** Absorption spectra of 20 µM cyt c in the presence of 5 µM p66Shc at 64-hour incubation time (red) or hypoxic condition (black) at 25 $^{o}$C.

**(C) Caspase 3 activity assay.** p66Shc and p46Shc effects on purified caspase 3 activity. Significance determined via t-tests. Shown as mean ± SD. N=5.

**(D) Alpha helix recovery after thermal melt.** p66Shc was maintained at 97 $^{o}$C for a minimum of 15 minutes, cooled overnight at room temperature (± CL), then tested on CD to determine % alpha helix recovery. Shown as mean ± SD. N=4.

**(E) p66Shc-mortalin MST binding results.** Shown as mean ± SD. N=4.

**(F) Shear effects on ROS activity.** Solutions were stirred (sheared) at 50% of max speed in a fluorimeter chamber 2 minutes prior to measuring ROS and during ROS measurements. Significance determined via t-tests, as indicated. Shown as mean ± SD. N=3.
